# Cholesterol and matrisome pathways dysregulated in human *APOE* ε4 glia

**DOI:** 10.1101/713362

**Authors:** Julia TCW, Shuang A. Liang, Lu Qian, Nina H Pipalia, Michael J. Chao, Yang Shi, Sarah E. Bertelsen, Manav Kapoor, Edoardo Marcora, Elizabeth Sikora, David M. Holtzman, Frederick R. Maxfield, Bin Zhang, Minghui Wang, Wayne W. Poon, Alison M. Goate

**Author notes:** Corresponding authors: Julia TCW and Alison Goate.

## Abstract

*Apolipoprotein E* (*APOE*) ε4 is the strongest genetic risk factor for Alzheimer’s disease (AD). Although its association with AD is well-established, the impact of *APOE* ε4 on human brain cell function remains unclear. Here we investigated the effects of *APOE* ε4 on several brain cell types derived from human induced pluripotent stem cells and human *APOE* targeted replacement mice. Gene set enrichment and pathway analyses of whole transcriptome profiles showed that *APOE* ε4 is associated with dysregulation of cholesterol homeostasis in human but not mouse astrocytes and microglia. Elevated matrisome signaling associated with chemotaxis, glial activation and lipid biosynthesis in *APOE* ε4 mixed neuron/astrocyte cultures parallels altered pathways uncovered in cell-type deconvoluted transcriptomic data from *APOE* ε4 glia and AD post-mortem brains. Experimental validation of the transcriptomic findings showed that isogenic *APOE* ε4 is associated with increased lysosomal cholesterol levels and decreased cholesterol efflux, demonstrating decoupled lipid metabolism. *APOE* ε4 glia also secrete higher levels of proinflammatory chemokines, cytokines and growth factors, indicative of glial activation. Thus, *APOE* ε4 induces human glia-specific dysregulation that may initiate AD risk.

---

Alzheimer’s disease (AD), the most common form of dementia, is characterized by widespread neurodegeneration, gliosis and two pathological hallmarks: amyloid-β (Aβ) plaques and tau-containing neurofibrillary tangles (NFTs)^1^. Familial, early-onset forms of AD are rare and caused by fully penetrant mutations in *APP* and *PSEN1/2*^2^. However, the majority of AD cases are sporadic, with an age at onset > 65yrs, and a complex genetic and environmental etiology^3^. Notably, the *Apolipoprotein E* (*APOE*) ε4 allele is the strongest genetic risk factor for AD; *APOE* ɛ4/ɛ4 (*APOE* 44) increases AD risk by 14-fold compared to *APOE* 33^4^ and is associated with increased AD pathology^5^. Although the association between *APOE* 4 and increased AD risk is well-established, the mechanisms underlying this effect in particular human brain cell types are not entirely clear. Prior studies in mouse models of AD have shown that *APOE* 4 increases Aβ seeding, decreases Aβ clearance, decreases synaptic plasticity and contributes to blood-brain barrier dysfunction^6,7^. Targeted replacement mice expressing human *APOE* (*APOE*-TR) 4 lack AD pathology with increasing age but exhibit *APOE* genotype-dependent effects on Aβ deposition and tau-dependent neurodegeneration when crossed with *APP* or *MAPT* mutant mouse models^8,9^.

While Apoe is normally expressed in astrocytes, recent studies have shown that it is highly upregulated in microglia in the context of the aged or diseased brain. This damage-associated microglial (DAM) polarization state is found near Aβ plaques in *APP* mice and is *Apoe*-dependent^10–12^. DAM exhibit downregulation of genes that are specifically expressed by microglia in the healthy brain (e.g., *P2ry12* and *Tgfbr1*) and upregulation of genes associated with lipid metabolism (e.g., *Apoe* and *Lpl*) and phagocytosis (e.g., *Axl* and *Trem2*). However, these studies were performed in the context of mouse Apoe or *Apoe* knockout (KO), which may differ from the effects of human *APOE* genotype^13^.

To more broadly address the mechanistic consequences of *APOE* 4 on human brain cell types, we employed human induced pluripotent stem cells (hiPSCs) generated from AD and control subjects of *APOE* 44 and *APOE* 33 genotypes. We examined effects of *APOE* genotype in microglia, astrocytes (hereinafter glia refers to both astrocytes and microglia) and brain microvascular endothelial cells (BMECs) as well as in mixed cultures of cortical neurons and astrocytes (hereinafter referred to as mixed cortical cultures). To compare the effects of *APOE* 4 in human and mouse cells, we purified primary microglia and astrocytes from *APOE*-TR and *Apoe* KO mice. Global transcriptomic profiling was carried out in each cell type to uncover gene expression changes associated with *APOE* 4 in human and mouse cells. Gene set enrichment and pathway analyses identified lipid metabolic deficits in *APOE* 4 hiPSC-glia and brain cell types. Further, cell-type deconvolution of global transcriptomic data from mixed cortical cultures and AD brain tissue homogenates revealed significant enrichment of matrisome pathways associated with chemotaxis, glial activation and lipid metabolism in *APOE* 4 glia. Finally, *in vitro* studies demonstrated decoupled cholesterol metabolism in isogenic *APOE* 44 astrocytes, leading to increased free cholesterol with decreased efflux, resulting in cholesterol accumulation in lysosomes. We further validated enhanced inflammatory chemokine and cytokine secretion from *APOE* 44 astrocytes. Taken together, our findings reveal *APOE* 4-driven molecular and cellular mechanisms that may contribute to AD risk.

## hiPSC-derived brain cell type differentiation

Thirteen hiPSC lines (derived from six and seven individuals with *APOE* 44 and 33 genotypes, respectively) were selected from 43 aged individuals of European ancestry, balanced for sex, disease status and controlled for AD genetic risk using a genetic risk score excluding *APOE* genotype (Fig. 1a, Extended Data Fig. 1a and Supplemental Data Table 1). After confirming the pluripotency and karyotype of the hiPSC lines (Extended Data Fig. 1c-d), each line was differentiated to four brain cell types: microglia, astrocytes, mixed cortical cultures and BMECs using published protocols^14–18^ (Fig. 1b). Mixed cortical cultures and astrocytes were differentiated from hiPSC-derived neural progenitor cells as described^14,15,18^. Mixed cortical cultures were primarily glutamatergic neurons with a minor portion of GABAergic, TH1+ dopaminergic neurons and astrocytes^14,18^ (Fig. 1e). Microglia were differentiated from a nearly pure population of CD43 positive hematopoietic progenitor cells (Extended Data Fig. 1e). BMECs were generated by exposing hiPSCs to endothelial growth factors. These differentiated cells were validated with cell type specific markers (Fig. 1e).

**Figure 1.**
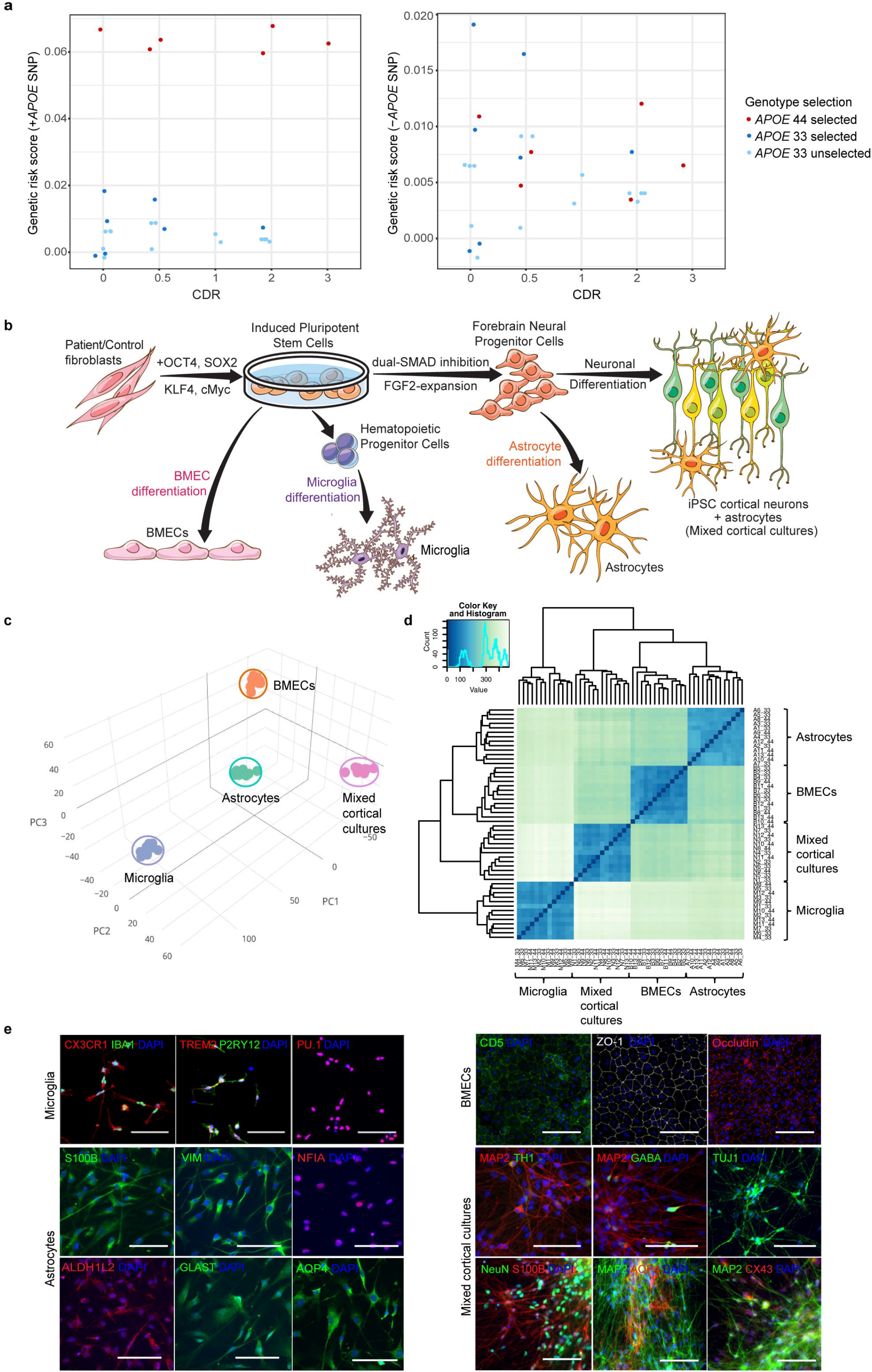
Genetic and *APOE* genotype based subject selection and brain cell type differentiation. **a.** Genetic risk score analysis with (left) or without *APOE* genotype (right) in *APOE* 33 and 44 lines (n = 13 selected lines out of total n = 43 screened lines) from multiple individual hiPSC lines. CDR, clinical dementia rating. **b.** Schematics of mixed cortical cultures, astrocytes, microglia and BMECs differentiation derived from *APOE* 33 and 44 hiPSCs. **c.** PCA of transcriptomic data from four cell types. **d.** Spearman correlation analysis of gene expression from 52 differentiated samples (n = 13 per cell type, n = 7 for *APOE* 33 and n = 6 for *APOE* 44 in each cell type). **e.** Representative immunofluoresent images of cell type specific markers for microglia (CX3CR1, IBA1, TREM2, P2RY12 and PU.1), astrocytes (S100B, VIM, NFIA, ALDH1L2, GLAST and AQP4), BMECs (CD5, ZO-1 and Occuludin), neurons (MAP2, TH1, GABA, TUJ1 and NeuN) with astrocytes (S100B, AQP4 and CX43) in mixed cortical cultures confirming cell type differentiations. Scale bar = 100μm.

To obtain comprehensive transcriptomic profiles for each of the four cell types, RNA sequencing (RNAseq) was performed. Principal component analysis (PCA) and Spearman correlation of the 52 transcriptomes (4 cell types x 13 lines) showed that samples clustered by cell type (Fig. 1c-d), suggesting that differentiation methods were robust and independent of *APOE* genotype. Each cell type expressed genes specific to and clustered with their corresponding primary human brain cell types (Extended Data Fig. 1f-g).

## Impaired lipid metabolism pathways in human *APOE* 44 glia

To investigate the effects of *APOE* 4, differentially expressed gene (DEG) analysis was performed in each hiPSC-derived brain cell type comparing *APOE* 44 to *APOE* 33. The number of DEGs in *APOE* 44 compared to *APOE* 33 was highest in astrocytes followed by mixed cortical cultures and microglia, no significant DEGs were observed in BMECs (Fig. 2a-b). To identify specific pathways affected by *APOE* 4, we performed fast preranked Gene Set Enrichment Analysis (fGSEA) using Molecular Signature Data Base (MSigDB) which includes canonical pathways, KEGG, Reactome, BioCarta and Gene Ontology (GO)-terms with FDR < 0.2 as a significance threshold^19^, following 1 million gene set permutations. Cholesterol biosynthesis was identified as the most significant and positively enriched gene set in *APOE* 44 microglia and astrocytes (Fig. 2c-d). Other lipid related gene sets such as Steroid biosynthesis (microglia and astrocytes) and Lipid and lipoprotein metabolism (astrocytes) were also enriched. In *APOE* 44 microglia, negatively enriched gene sets included HDL-mediated lipid transport, Generic transcription and Lysosome.

**Figure 2.**
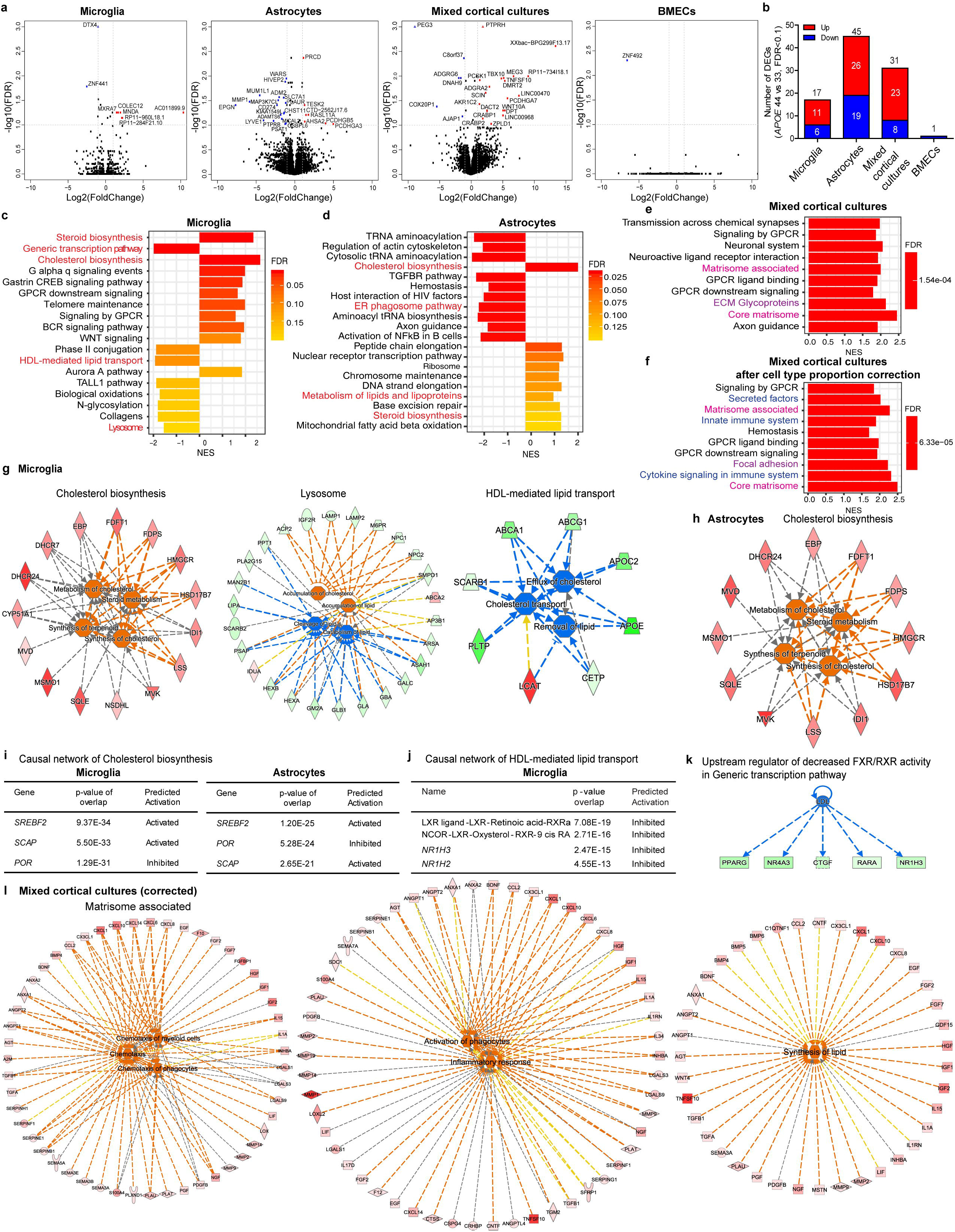
Enriched Lipid- and Matrisome-related pathways associated with *APOE* 44 in hiPSC-derived brain cell types. **a.**Volcano plot comparison of *APOE* 44 versus *APOE* 33 in microglia, astrocytes, mixed cortical cultures and BMECs. Average log2 (fold change) by log10 (FDR) is shown for all genes. Genes upregulated (red) and downregulated (blue) by 2-fold change with FDR < 0.1 are labeled with gene names. **b.** The number of significantly and differentially expressed genes in **a**. **c-f.** Top significant pathways from gene set enrichment analysis of DEGs of *APOE* 44 compared to *APOE* 33 in microglia (**c**), astrocytes (**d**), mixed cortical cultures (positive enrichment) (**e**) and mixed cortical cultures after cell type proportion correction (positive enrichment) (**f**). Color text label of red, lipid-related; magenta, matrisome-related; purple, ECM-related; blue, immune-related pathways. **g.** Functional pathway analysis of Cholesterol biosynthesis, Lysosome and HDL-mediated lipid transport pathways enriched in *APOE* 44 microglia from fGSEA. Red and orange color are upregulated genes or functions, and green or blue colors are downregulated genes or functions. **h.** Functional pathway analysis of Cholesterol biosynthesis, enriched in *APOE* 44 astrocytes from fGSEA. **i.** Causal network analysis of Cholesterol biosynthesis in *APOE* 44 microglia and astrocytes. **j.** Causal network analysis of HDL-mediated lipid transport in *APOE* 44 microglia. **k.** Upstream regulator of downregulated FXR/RXR activity from top canonical pathways of Generic transcription pathway, negatively enriched in *APOE* 44 microglia. **l.** Functional pathway analysis by ingenuity pathway of Matrisome associated in *APOE* 44 vs *APOE* 33 in cell type proportion corrected mixed cortical cultures.

To determine which molecular and cellular functions, upstream regulators and causal networks of each enriched gene set are altered in *APOE* 44 compared to *APOE* 33 glia, we performed causal network analysis of fGSEA-enriched gene sets using Ingenuity Pathway Analysis (IPA)^20^. DEGs of the gene set enriched in Cholesterol biosynthesis is predicted to increase synthesis of cholesterol and terpenoids and upregulate cholesterol and steroid metabolism in *APOE* 44 compared to *APOE* 33 microglia (Fig. 2g). DEGs of the gene set enriched in Lysosome is predicted to enhance accumulation of cholesterol and decrease lipid clearance and catabolism (Fig. 2g). Reduced cholesterol efflux is predicted by a gene set enriched in HDL-mediated lipid transport in *APOE* 44 microglia (Fig. 2g). Likewise, *APOE* 44 astrocytes show upregulation of cholesterol synthesis and metabolism (Fig. 2h). Causal network analysis showed that upstream regulators (e.g. transcription factors regulating cholesterol synthesis^21,22^), *SCAP* and *SREBF2* are significantly activated but *POR* (Cytochrome P450 oxidoreductase) is inhibited in both *APOE* 44 microglia and astrocytes (Fig. 2i). In contrast, the *LXR/RXR*, lipid-responsive transcription factors that regulate *APOE* expression and cholesterol efflux, *LXR* ligands and the *NR1H2* and *NR1H3* nuclear receptors, are predicted to be downregulated (Fig. 2j), suggesting that lipid/cholesterol metabolism and transport may be altered in astrocytes and microglia as a consequence of *APOE* 44 genotype. The most negatively enriched, Generic transcription pathway in *APOE* 44 microglia involves inhibition of *FXR/RXR* activity (Fig. 2c and Extended Data Fig. 2a). The causal network of Generic transcription pathway predicts FXR/RXR signaling will be downregulated by decreased low-density lipoprotein (LDL), suggesting suppression of the lipid efflux system due to lower intracellular cholesterols in *APOE* 44 microglia (Fig. 2k). Thus, global transcriptomic analyses reveal that *APOE* 44 genotype is associated with higher cholesterol synthesis and lower catabolism/efflux, suggesting either that cholesterol levels are lower in *APOE* 44 cells or that the cholesterol sensing mechanism is impaired in *APOE* 44 microglia and astrocytes.

In addition, Generic transcriptional pathway in *APOE* 44 microglia is associated with cell cycle downregulation, leading to decreased cell proliferation and RNA transactivation (Extended Data Fig. 2a-b). In *APOE* 44 astrocytes one of the most negatively enriched gene sets is regulation of the actin cytoskeleton (Fig. 2d). Interestingly, IPA of this gene set highlights decreased mitosis, mitogenesis and cell cycle progression (proliferation) and increased cell death regulating both apoptosis and necrosis (Extended Data Fig. 2c-d). It further predicts reduced astrocyte projections, suppressed endocytosis/phagocytosis and decreased cellular homeostasis in *APOE* 44 glia (Extended Data Fig. 2d).

## Glial origin matrisome gene set/pathway enriched in *APOE* 44 cells and AD brains

Examination of mixed cortical cultures identified a number of significant DEGs in *APOE* 44 compared to *APOE* 33 (Fig. 2a-b). Given the heterogeneous nature of these cultures we first measured the proportions of neurons and glia using four different algorithms: digital sorting algorithm (DSA)^23^, population-specific expression analysis (PSEA)^24^, non-negative matrix factorization (ssKL)^25^, and a PCA-based method modified from CellCODE (BRETIGEA, BRain cEll Type specIfic Gene Expression Analysis)^26^ (Extended Data Fig. 2e-f). Although the majority of cells were neurons, all cultures contained 5-25% astrocytes (Extended Data Fig. 2f). Before and after astrocyte proportion correction in the mixed cortical cultures by DSA, the best fit algorithm validated on primary human brain cell types (Extended Data Fig. 3b), we found consistent enrichment of matrisome-related pathways including Matrisome associated, Core matrisome and Extracellular matrix (ECM) glycoproteins (Fig. 2e-f). After the correction, we also observed significant enrichment of innate immune/cytokine pathways (Fig. 2f). The *in silico* ‘Matrisome’ is defined as the ensemble of ECM proteins and associated factors that are built by compiling proteomics data on the ECM composition^27,28^. ECM proteins provide biochemical cues interpreted by cell surface receptors like integrins and initiate signaling cascades governing cell survival, proliferation and differentiation^29,30^. While the ‘Core matrisome’ comprises ECM glycoproteins, collagens and proteoglycans, ‘Matrisome associated’ proteins include ECM-affiliated proteins, ECM regulators and secreted factors including growth factors and cytokines released from both neurons and glia^27^. Further analysis of Matrisome associated pathway in *APOE* 44 mixed cortical cultures showed three functional modules: upregulated chemotaxis of phagocytes/myeloid cells, activation and inflammatory responses of cells and synthesis of lipids (Fig. 2l). Additionally, Core matrisome pathway in *APOE* 44 cells displays increased levels of ECM molecules that support cell attachment and migration (Extended Data Fig. 2g). The presence of matrisome-related pathways in *APOE* 44 mixed cortical cultures but not in the pure astrocyte cultures led us to hypothesize that enriched matrisome is associated with communication between neurons and astrocytes^31^ and may also be observed in *APOE* 44 brains.

To determine whether matrisome pathways are also upregulated in human *APOE* 44 or AD brains, we performed DEG analyses in RNAseq data from the MSBB and ROSMAP cohorts comparing *APOE* 44 versus (vs) *APOE* 33 in deconvoluted cell type-specific whole transcriptome profiles of AD brains as well as AD vs control in *APOE* 33 carriers. Using the cell type proportion estimates, DEG statistics of *APOE* 44 compared to *APOE* 33 in each cell type were computed, following deconvolution for each cell type in each brain region using the csSAM algorithm^32^ (Extended Data Fig. 3a-c). Matrisome-related signals were observed in *APOE* 44 astrocytes from superior temporal gyrus (STG, BA22), parahippocampal gyrus (PHG, BA36) and inferior frontal gyrus (IFG, BA44) in MSBB and dorsolateral prefrontal cortex in ROSMAP from AD brains (Fig. 3a-e). Although *APOE* 44 microglia do not exhibit enriched matrisome pathways, they do show positive enrichment of ECM, chemokine, innate immune and proinflammatory cytokine signaling pathways in all brain regions examined (Fig. 3f-i). Next, we tested whether matrisome-related pathways are associated with AD, independent of *APOE* genotype. Pathway analysis of the brain data within *APOE* 33 genotype identified that Matrisome and Core matrisome are the most significantly upregulated pathways in AD brains diagnosed by various traits compared to controls (Fig. 3j). In addition, cell adhesion pathways (ECM/integrin) and inflammatory pathways (cytokine pathways) are positively enriched in AD brains (Fig. 3j). Thus, matrisome pathways were strongly enriched in glial cells of *APOE* 44 carriers and in AD regardless of *APOE* genotype. Further pathway analysis of DEGs from AD vs control within *APOE* 33 individuals showed similar modules to those identified in *APOE* 44 vs *APOE* 33 mixed cortical cultures (Fig. 2l and Extended Data Fig. 3n), indicating that the transcriptomic profile of *APOE* 44 mixed cortical cultures resembles a phenotype detected in AD compared to control brains regardless of *APOE* genotype. In addition, combining *APOE* 44 genotype and AD phenotype effect, we observed significantly enriched matrisome in glia deconvoluted from all brain regions and AD brains (Extended Data Fig. 3d-m).

**Figure 3.**
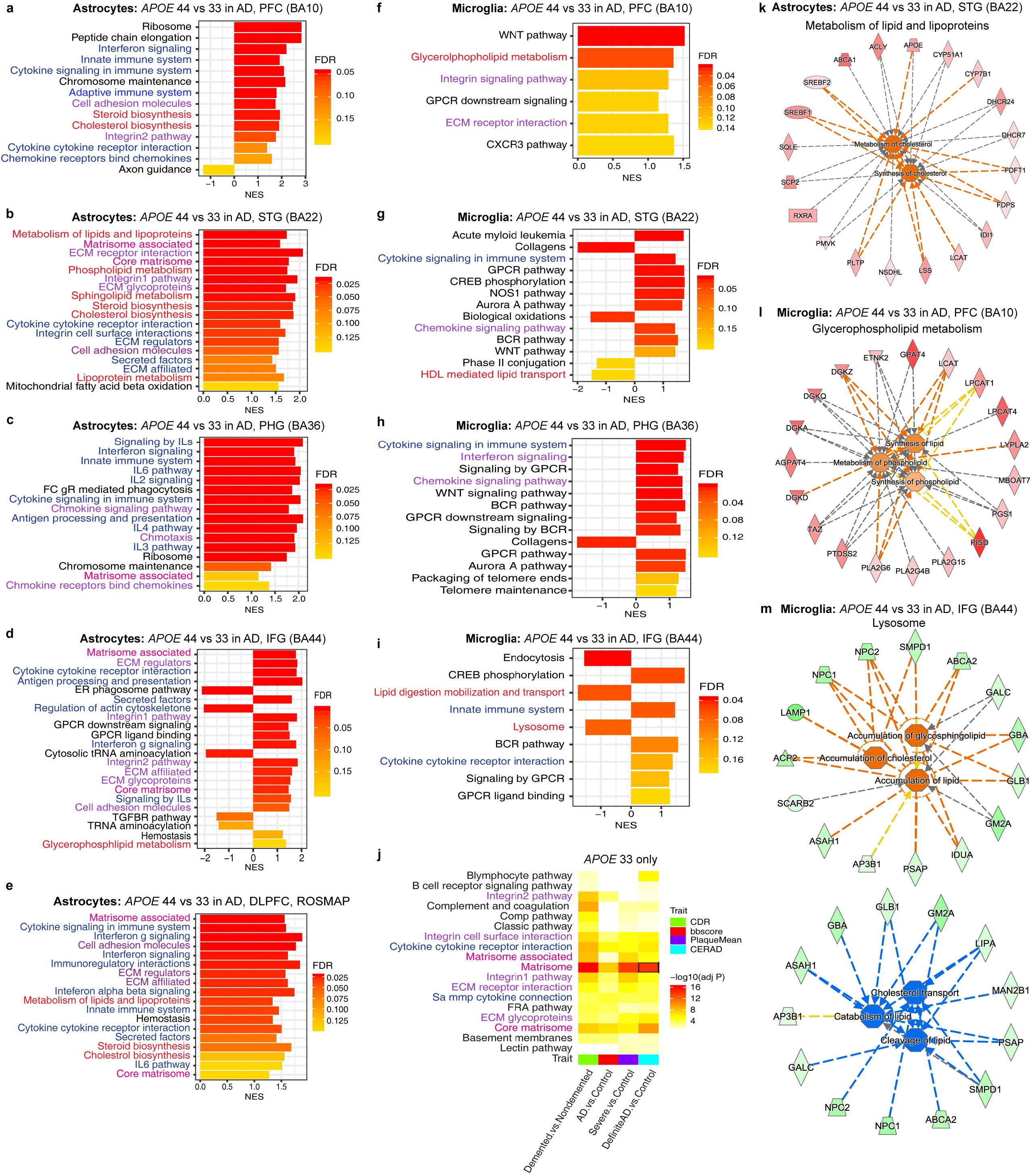
Enrichment of matrisome in *APOE* 44 hiPSC-mixed cortical cultures, deconvoluted *APOE* 44 glia and AD brains in global transcriptomic data. **a-i.** fGSEA of DEGs of *APOE* 44 compared to *APOE* 33 in each cell type (astrocyte or microglia) after cell type deconvolution in various regions of AD brain from multiple cohorts (MSBB and ROSMAP). PFC, prefrontal cortex; STG, superior temporal gyrus; PHG, parahippocampal gyrus; IFG, inferior frontal gyrus; DLPFC, dorsolateral prefrontal cortex. **j.** Upregulated canonical pathways of DEGs in different regions of brain comparing various criteria of AD-related phenotypes only in *APOE* 33 carriers. Functional pathways of Matrisome (marked with a black box) was analyzed in **Extended Data Fig. 3n**. **k-l.** Functional pathway analysis of lipid-related pathways in *APOE* 44 astrocytes in STD (BA22) (**k**) and microglia in PFC (BA10) and IFG (BA44) (**l-m**) of MSBB.

We also investigated whether pathways enriched in *APOE* 44 hiPSC-astrocytes and microglia can be identified in *APOE* 44 human brains (AD or control) in a cell type-specific manner. *APOE* 44 astrocytes- and microglia-specific whole transcriptome profiles deconvoluted from AD brains showed positive enrichment of Cholesterol/Steroid biosynthesis in astrocytes of frontal cortex (BA10 and ROSMAP) and STG and additional enrichment of Sphingolipid, Phospholipid metabolism in astrocytes of STG and IFG and microglia of PFC (Fig. 3a-i and Extended Data Fig. 3d-e, 3h-l). It demonstrates that one of the major gene sets enriched in *APOE* 44 hiPSC-glia is also enriched in *APOE* 44 brain-transcriptome-deconvoluted glia. The negatively enriched HDL-mediated lipid transport and Lysosome in *APOE* 44 hiPSC-microglia was also observed in *APOE* 44 AD microglia of IFG and STG (Fig. 3i). Further functional predictions reveal that positively enriched gene sets associated with several lipid metabolism pathways in *APOE* 44 astrocytes and microglia lead to increased synthesis of cholesterol, similar to *APOE* 44 hiPSC-glia (Fig. 3k-l). Reduction of lysosomal gene expression in *APOE* 44 microglia leads to a predicted accumulation of cholesterol and other lipids likely due to decreased catabolism of lipid and cholesterol transport (Fig. 3m). Thus, dysregulated transcriptome of lipid metabolism in *APOE* 44 hiPSC-microglia and cholesterol/steroid synthesis in *APOE* 44 hiPSC-astrocytes recapitulates changes observed in *APOE* 44 brain-transcriptome-deconvoluted glia.

Together, our cell type proportion correction, followed by deconvolution of the transcriptome reveal that *APOE* 44 hiPSC-mixed cortical cultures resemble the transcriptome observed in human AD brains and that the significantly enriched gene sets in both conditions are associated with the support of glial migration, activation and synthesis of lipid, which are derived from *APOE* 44 glia as a consequence of communication between neurons and glia.

## Human specific effects of *APOE* 44 on glial transcriptomes

Targeted replacement of the endogenous murine *Apoe* gene with human *APOE* has been utilized to understand *APOE* 4 risk in the background of *APP* or *MAPT* mutations^9,33^. To unravel mouse glial cell type-specific effects of *APOE* 4, microglia and astrocytes were purified from 16 mouse fetal brains of *APOE* 44 (n=6), *APOE* 33 (n=6) and *Apoe* KO (n=4) mice. Cell type specific markers were used to confirm purity of the cultures (Fig. 4a). Clustering analysis of 32 samples (n=16 for each cell type) based on Spearman correlation revealed that they are well-clustered by cell type (Fig. 4b). PCA analysis shows that *APOE* 33 and *APOE* 44 cells cluster together within each cell type, while *Apoe* KO microglia and astrocytes are well-separated from both *APOE* 33 and *APOE* 44 cells (Fig. 4c). Consistent with human glia, mouse astrocytes have a higher number of DEGs than microglia in *APOE* 44 vs *APOE* 33, possibly because *APOE* is more highly expressed in astrocytes than microglia under the baseline condition used in our experiments (Fig. 4d and Extended Data Fig. 4a). After homology conversion of mouse to human DEGs, followed by fGSEA Extended Data Fig. 4c), we observed that Matrisome associated, ECM affiliated and Interferon/Cytokine/Adaptive immune pathways are enriched in both *APOE* 44 mouse microglia and astrocytes (Fig. 4f). Of note, functional analyses of Matrisome associated pathway in *APOE* 44 mouse microglia and astrocytes do not exhibit lipid-related dysregulation in contrast to human cells (Fig. 4i vs Fig. 2l). This suggests that although *APOE* 44 mouse glia resemble *APOE* 44 human glia for enrichment of matrisome and inflammation pathways, lipid metabolic dysregulation appears to be specific to human *APOE* 44 glia.

**Figure 4.**
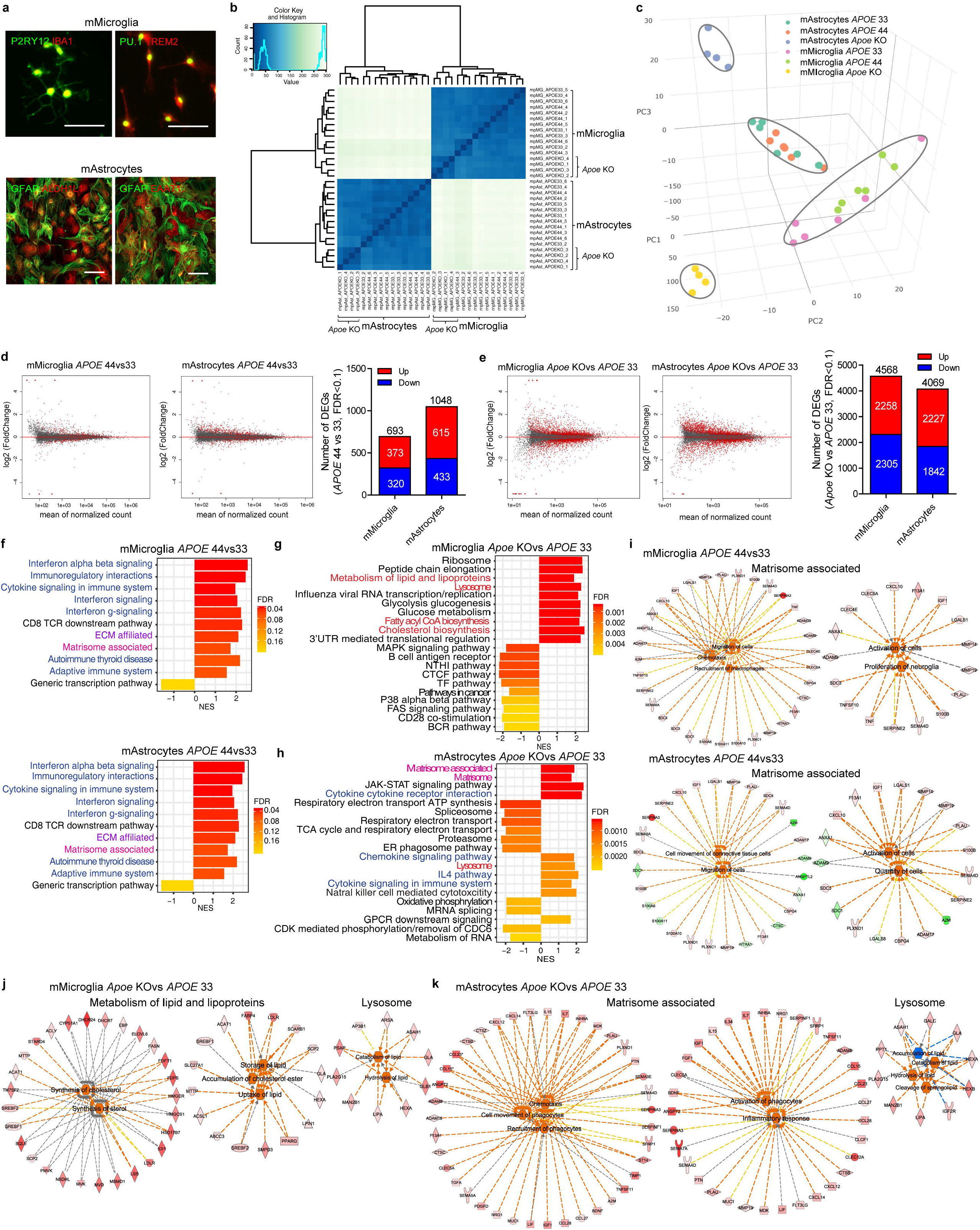
Shared Matrisome pathways in human and mouse *APOE* 44 glia while human glia specific lipid metabolic dysfunction, which resemble mouse *Apoe* KO but not mouse *APOE* 44 glia. **a.** Immunocytochemistry for cell type specific markers on purified mouse microglia (mMicroglia) (P2RY12, IBA1, PU.1 and TREM2) and astrocytes (mAstrocytes) (GFAP, ALDH1L1 and EAAT1). Scale bar = 100μm. **b.** Spearman correlation analysis of transcriptomic data from 3 genotypes and 2 cell types (n = 16 per cell type, n = 6 for *APOE* 33 and *APOE* 44 and n = 4 for *Apoe* KO in each cell type). **c.** PCA of transcriptomic data from 3 genotypes and 2 cell types. **d.** MA (M: log ratio and A: mean average) plots of *APOE* 33 vs *APOE* 44 comparisons in mMicroglia and mAstrocytes (left). Average log2 (fold change) by mean of normalized count is shown for all genes. Genes marked red are FDR < 0.1. The number of significantly and differentially expressed genes in MA plots (right) **e.** MA plots of *Apoe* KO vs *APOE* 33 comparisons in in mMicroglia and mAstrocytes (left) and the number of significant DEGs in MA plots (right). **f-h.** fGSEA of DEGs of *APOE* 44 (**f**) or *Apoe* KO (**g-h**) compared to *APOE* 33 in mMicroglia and mAstrocytes. **i.** Ingenuity pathway analysis of Matrisome associated, significantly enriched pathway in *APOE* 44 mMicroglia (top) and mAstrocytes (bottom). **j-k.** Ingenuity pathway analysis of Metabolism of lipid and lipoprotein and Matrisome associated, enriched in *Apoe* KO mMicroglia and mAstrocytes, respectively and lysosome in *Apoe* KO of both cell types compared to *APOE* 33.

*Apoe* KO exhibits significantly higher number of DEGs (7-9 fold and 4-7 fold) compared to *APOE* 33 or *APOE* 44 microglia and astrocytes, respectively (Fig. 4e and Extended Data Fig. 4a-b). Further assessment of *Apoe* KO compared to *APOE* 33 in mouse microglia and astrocytes revealed cell type-specific effects. *Apoe* KO microglia but not astrocytes display enriched Cholesterol biosynthesis and other lipid regulatory pathways (Fig. 4g-h) and further confirm *Apoe* loss-of-function in regulation of lipid synthesis (Fig. 4j). In *Apoe* KO astrocytes, matrisome-related and immune pathways are positively enriched (Fig. 4h) but not lipid regulatory functions (Fig. 4k). Thus, upregulated lipid metabolism is specific to mouse *Apoe* KO (i.e. absent from *APOE* 44) microglia, similar to observations in human *APOE* 44 glia. However, in contrast to human *APOE* 44 microglia, the Lysosome gene set displays positive enrichment with upregulation of lipid catabolism in mouse *Apoe* KO microglia and astrocytes (Fig. 4j-k). Further analysis of *Apoe* KO vs *APOE* 44 DEGs shows consistent results to *Apoe* KO vs *APOE* 33 DEGs, demonstrating that *Apoe* KO-driven dysregulation is distinct from *APOE* 44 dysregulation in mouse (Extended Data Fig. 4d-e). Together, these analyses demonstrate that mouse *APOE* 44 glia partially recapitulate ECM and immune signals but not lipid metabolism dysregulation observed in human *APOE* 44 glia. Mouse *Apoe* KO shows distinct transcriptomic dysregulation characterized by upregulation of lysosomal lipid catabolism in both microglia and astrocytes, upregulation of cholesterol synthesis in microglia and upregulation of matrisome pathway in astrocytes.

## Elevated cholesterol in *APOE* 44 hiPSC- astrocytes due to decoupled lipid metabolism

Although we saw consistent differences in the transcriptomic signatures from *APOE* 44 vs *APOE* 33 hiPSC-derived neural cultures, there is extensive linkage disequilibrium around the *APOE* locus, and thus the observed differences cannot be unambiguously attributed solely to *APOE* genotype. To demonstrate that the changes in cholesterol metabolism, predicted from the global transcriptomic analyses are indeed caused by *APOE* genotype, we created 12 isogenic *APOE* hiPSC lines from two *APOE* 44 individuals using CRISPR/Cas9 gene-editing (Extended Data Fig. 5a-b). Potential off-targets (quality score ≥ 0.5) using the designed gRNA were confirmed negative, all isogenic lines were karyotypically normal, and their genetic identities matched the original fibroblasts (Supplemental Data Table 6).

Since transcriptomic analysis identified pathways consistent with lipid accumulation in human *APOE* 44 glia, we examined sterol metabolism *in vitro*. Cellular cholesterol levels measured by Gas Chromatography coupled to Mass Spectrometry (GC-MS) showed 20% increase in total cholesterol and a similar increase in free (or unesterified) cholesterol but not cholesteryl ester levels in *APOE* 44 compared to isogenic *APOE* 33 astrocytes (Fig. 5a), indicating that the increased total cholesterol represents free cholesterol. Consistently, we observed increased filipin levels in *APOE* 44 compared to *APOE* 33 astrocytes, indicative of elevated free cholesterol in *APOE* 44 astrocytes (Fig. 5b-c) and elevated levels of HMG-CoA reductase (HMGCR), the enzyme responsible for the rate limiting step in cholesterol biosynthesis, in *APOE* 44 compared to *APOE* 33 astrocytes (Fig. 5d-e). To assess cholesterol accumulation in the presence of excess extracellular lipid, purified LDL particles were added to astrocyte media. Intracellular free cholesterol levels were higher in *APOE* 44 compared to *APOE* 33 astrocytes at baseline and after LDL challenge (Fig. 5b-c), indicating intracellular lipid dysregulation in *APOE* 44 glia. The average filipin intensity measured per cell by filipin assay is in agreement with differences in free cholesterol measured by GC-MS. The intracellular localization of free cholesterol was observed by co-labeling the astrocytes with filipin and endocytosed FITC-Dextran, a marker for lysosomes. The co-localization of filipin with FITC-Dextran appears as yellow puncta, clearly indicating that free cholesterol is significantly higher in *APOE* 44 astrocytes and that much of the additional cholesterol is in lysosomes (Fig. 5f). This phenotype appears to be cell type-specific since the parental fibroblasts do not show a genotype-dependent difference in cholesterol levels (Extended Data Fig. 5d-e). These findings demonstrate defective cholesterol accumulation in *APOE* 44 glia.

**Figure 5.**
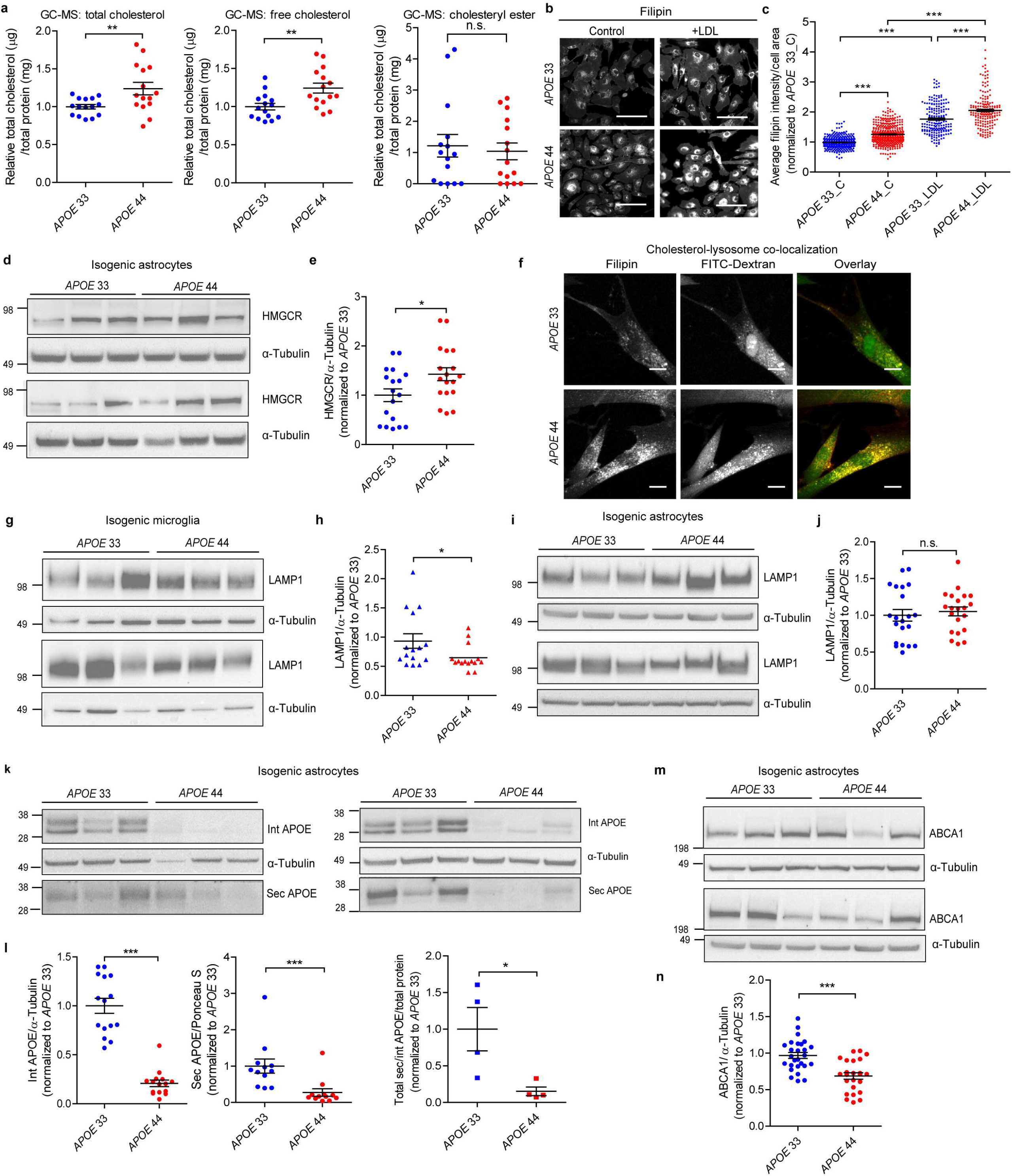
Isogenic *APOE* 44 glia accumulate intracellular cholesterol due to decoupled lipid metabolism. **a.** Level of total cholesterol, free cholesterol and cholesteryl ester per milligram total protein in isogenic *APOE* 33 and *APOE* 44 astrocytes measured by Gas Chromatography and Mass Spectrometry (GC-MS) (n = 3 isogenic lines per *APOE* genotype, 4 independent experiments with 3 replicates). **b.** Representative fluorescence microscopy images of filipin staining in isogenic *APOE* 33 and *APOE* 44 astrocytes with or without LDL treatment. Scale bar = 100μm. **c.** Quantification of whole fields of filipin images of isogenic *APOE* 33 and *APOE* 44 astrocytes (n = 3 isogenic lines per genotype, 5 independent experiments with ~ 20 quantified areas per experiment, each dot represents average intensity per cell area): average filipin intensity by cell area normalized to *APOE* 33. C, no serum control. **d-e.** Representative immunoblot images (**d**) and quantification (**e**) of HMGCR in isogenic *APOE* astrocytes (n = 6 isogenic lines per *APOE* genotype, 3 independent experiments). **f.** Representative sum projection of confocal images of filipin (red) and endocytosed FITC-Dextran (green) in *APOE* 33 and *APOE* 44 astrocytes. Yellow puncta in overlay images indicate lysosomal localization of cholesterol. Size bar = 15μm**. g-j.** Representative immunoblot images and quantification of LAMP1 in isogenic *APOE* 33 and *APOE* 44 microglia (**g-h**) and astrocytes (**i-j**). **k-l.** Representative immunoblot images (**k**) and quantification (**l**) of intracellular and secreted APOE and their ratios in isogenic *APOE* 33 and *APOE* 44 astrocytes. Raw images for supernatant calculations are in **Supplementary fig. 1. m-n.** Representative immunoblot images (**m**) and quantification (**n**) of ABCA1 in isogenic *APOE* astrocytes. Each column in immunoblot images represents an independent CRISPR line per genotype. One-way unpaired t-test for genotype comparisons and One-way ANOVA with Bonferroni post-corrections for comparisons of multiple treatments. *, p < 0.05, **, p < 0.01, ***, p < 0.001.

Next, we examined proteins involved in endo/lysosomal organelles and lipid efflux mechanisms since the transcriptomic data predicts deficits in lysosomal catabolism as well as cholesterol efflux. Consistent with the transcriptomic data showing microglial specific downregulation of genes in the Lysosome gene set including *LAMP1*, LAMP1 was significantly decreased in *APOE* 44 microglia but unchanged in astrocytes (Fig. 5g-j). Deficiency of LAMP1, which resides in lysosomal/late endosomal membranes and directly binds cholesterol, is associated with a defect in transport of cholesterol to the site of esterification in the endoplasmic reticulum, resulting in intracellular cholesterol accumulation^34,35^. Of note, *APOE* 44 astrocytes have much lower levels of both intracellular (80% reduction) and secreted (63% reduction) APOE compared to *APOE* 33 astrocytes (Fig. 5f-g). The ratio of secreted to intracellular APOE was also lower in *APOE* 44 cells (Fig. 5k-l), suggesting decreased efflux of lipids. ATP-binding cassette transporter ABCA1, a major regulator of cellular cholesterol homeostasis through the transport of lipids via APOE, was also significantly decreased in *APOE* 44 compared to *APOE* 33 astrocytes (Fig. 5m-n).

Taken together, isogenic *APOE* 44 astrocytes accumulate intracellular free cholesterol but exhibit decreased levels of lipid carriers and transporters with unchanged or lower levels of lysosomal lipid carriers. Combined with the transcriptomic data these results indicate dysregulation of lipid/cholesterol metabolism in *APOE* 44 glia. Specifically, lipid/cholesterol biosynthesis is elevated and lipid/cholesterol catabolism is decreased in cells that exhibit elevated cholesterol levels, suggesting that cholesterol flux through the lysosome to the endoplasmic reticulum is impaired. In the presence of cholesterol overload, *APOE* 44 cells take up and accumulate more lipid than *APOE* 33 cells, even though baseline levels of lipid are already higher. Together, these results suggest that cholesterol metabolism is dysregulated in *APOE* 44 glia.

## Increased chemokines/cytokines profile of isogenic *APOE* 44 astrocytes

Cell type-deconvolution analysis of AD brain transcriptomes showed a significant enrichment of matrisome-related pathways in *APOE* 44 astrocytes. To examine this pathway *in vitro* we measured a panel of secreted proteins that include chemokines, cytokines and growth factors using a Luminex multiplex immunoassay in isogenic human *APOE* astrocytes. Hierarchical clustering of the data showed that samples are clustered by *APOE* genotype rather than individual (Extended Data Fig. 6a). Nearly half of the proteins were differentially expressed by *APOE* genotype (24 of 45). The top 12 differentially secreted proteins by *APOE* genotype include the chemokines (SDF-1a (CXCL12), Gro-alpha/KC (CXCL1), MIP-1b (CCL4), Eotaxin (CCL11), IP-10 (CXCL10) and RANTES (CCL5)), cytokines (IL-8, LIF and IL-6) and growth factors (VEGF-A, HGF and VEGF-D) (Fig. 6a and Extended Data Fig. 6b). Spearman correlation coefficient analysis of these 12 proteins showed a stronger correlation between these markers in *APOE* 33 vs *APOE* 44 astrocytes (Extended Data Fig. 6c-e). When absolute protein levels were compared between *APOE* 33 and *APOE* 44 astrocytes, 10 of these proteins showed significantly higher levels in the media from *APOE* 44 glia, supporting elevated chemotactic molecules and cytokines in *APOE* 44 astrocyte transcriptomes.

**Figure 6.**
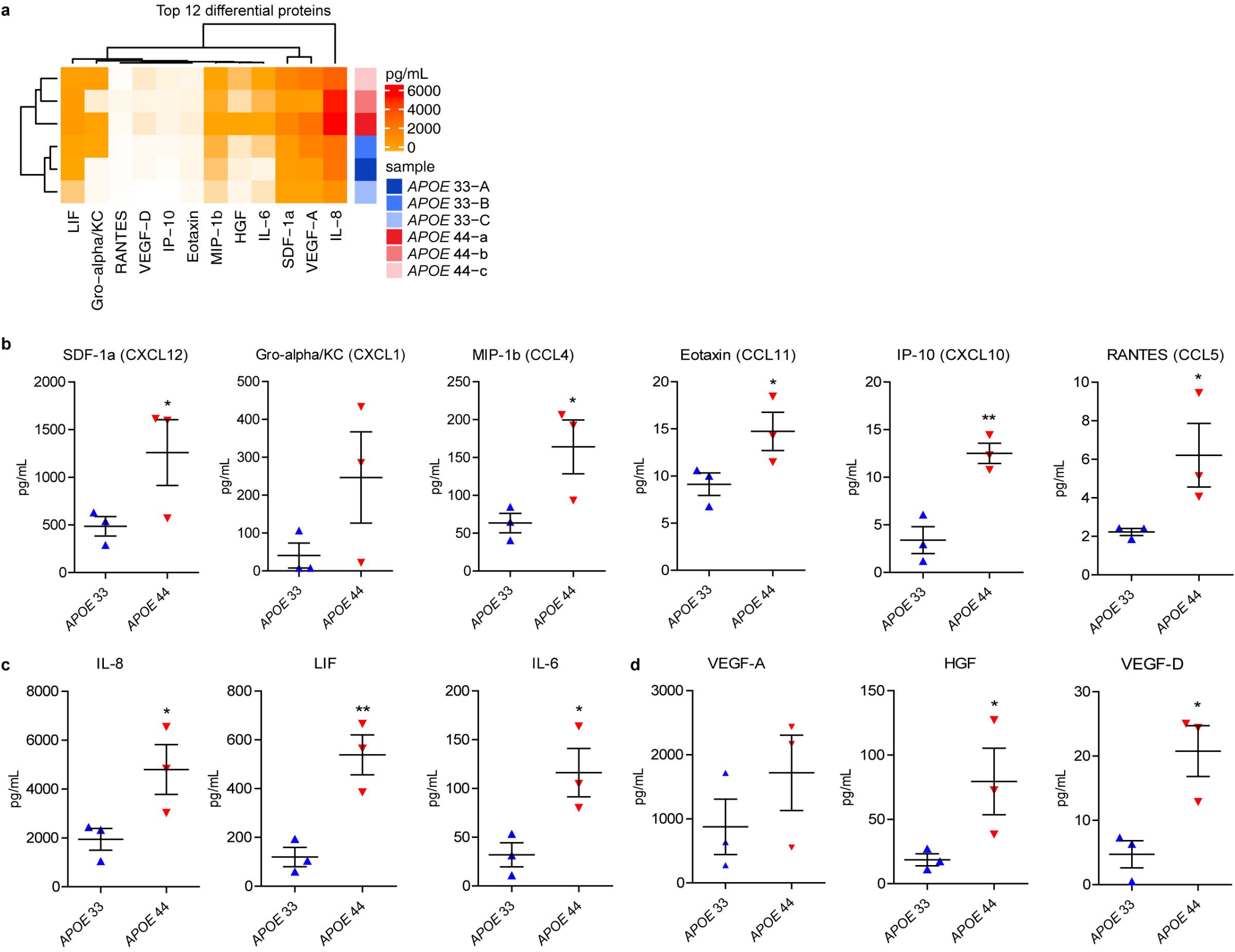
Secreted proteins from isogenic *APOE* 44 astrocytes show increased profiles of chemokines and cytokines compared to *APOE* 33 cells. **a.**Clustering heatmap for top 12 secreted proteins by *APOE* genotype screened from 45-plex human panel 1 including chemokines, cytokines and growth factors. A-C and a-c are independent genome edited CRISPR lines of isogenic *APOE* 33 and *APOE* 44 astrocytes. **b-d.** Quantification of chemokines (b), cytokines (c) and growth factors (**d**) (mean ± SEM) secreted by isogenic *APOE* 33 and *APOE* 44 lines (n = 3 isogenic lines per *APOE* genotype, one dot represents 2 experiments) from the same number of seeded cells. One-way unpaired t-test for genotype comparisons. *, p < 0.05, **, p < 0.01.

## Concluding remarks

An *Apoe*-GFP reporter study in mice demonstrated that Apoe is expressed in astrocytes and small subsets of microglia, neurons and endothelial cells at baseline but is increased in microglia and neurons after kainic acid exposure^36^. Single cell RNAseq studies have demonstrated that *Apoe* is one of the most significantly upregulated genes in a subset of microglia, termed DAM^11^. In mouse models of amyloidosis, these microglia are specifically located around amyloid plaques^12^. KO of *Apoe* in microglia blocks this polarization in response to damage, indicating that Apoe plays a critical role in these cells, which are characterized at the transcriptomic level by upregulation of phagocytosis and lipid metabolism genes and downregulation of homeostatic genes^10^. In this study, we have used global transcriptomics and *in vitro* functional studies to determine the impact of human *APOE* 44 genotype on gene expression and function in four human brain cell types: microglia, astrocytes, mixed cortical cultures and BMECs derived from 13 hiPSC lines of different *APOE* genotypes and 12 isogenic lines. Human *APOE* 44 microglia and astrocytes show dysregulation of cholesterol metabolism compared to *APOE* 33 cells. Specifically, *APOE* 44 microglia display increased lipid biosynthesis, lipid accumulation and decreased lipid catabolism and efflux, suggesting a decoupling of lipid synthesis and catabolism. Similar pathway dysfunction was observed in *APOE* 44 glia deconvoluted from human brains derived from multiple AD cohorts.

In *APOE* 44 hiPSC-mixed cortical cultures compared to *APOE* 33, matrisome related, ECM, immune pathways were enriched in fGSEA, which replicated findings from brain-transcriptome-deconvoluted *APOE* 44 glia and AD brains. Upregulation of ECM caused by astrogliosis in brain is associated with accumulation of amyloid plaques in AD and the formation of the glial scar after CNS injuries, which establishes a mechanical barrier inhibiting neurite outgrowth^37,38^. Thus, the hiPSC-mixed cortical model reveals that the *APOE* 44 mixed neuronal and glial condition recapitulates some aspects of the AD brain environment and suggests a reactive astrocyte state that upregulates the secretion of ECM, cytokines and growth factors. The DEGs of the Matrisome associated gene set enriched in *APOE* 44 mixed cortical cultures also showed increased synthesis of lipids that were discovered in pure populations of astrocytes, indicating a link between inflammatory response and lipid synthesis. Comparison of mouse and human glial transcriptomes indicates that mouse *APOE* 44 glia only partially capture the defects observed in human *APOE* 44 glia: Matrisome associated, ECM and inflammatory pathways are enriched in *APOE* 44 mouse microglia and astrocytes but not dysfunction of lipid metabolism pathways, stressing the importance of studying *APOE* genotype-dependent effects in human model systems. Interestingly, *Apoe* KO transcriptomes from both microglia and astrocytes were strikingly different from both *APOE* 33 and *APOE* 44, which were much more similar to each other. Despite this, DEG analysis of *Apoe* KO vs *APOE* 33 and *APOE* 44 identified many of the same dysregulated pathways seen between *APOE* 33 and *APOE* 44 glia.

Niemann-Pick Type C (NPC) is an inherited neurodegenerative disease caused by mutations in either *NPC1* or *NPC2* that results from a failure of endo/lysosomal cholesterol trafficking, causing lipid accumulation in lysosomes^39^. Although NPC (sometimes referred to as “Childhood Alzheimer’s”) differs in major respects from AD^40,41^, our observation of intracellular cholesterol accumulation, particularly in the lysosomes, coupled with decreased lipid efflux in *APOE* 44 glia, shares similarities with the molecular consequences of *NPC* loss-of-function, and may represent one of the earliest molecular changes leading to lipidosis in AD^42^.

In summary, we demonstrate human-specific and brain cell type-dependent transcriptional and cellular effects of *APOE* 44 in hiPSC-derived cultures. Furthermore, these changes mimic *APOE* 44-dependent transcriptomic changes detected in AD brain and uncover deficits in lipid homeostasis and glial activation. These studies suggest that therapeutic approaches aimed at restoring lipid homeostasis in glia may be beneficial in AD, particularly in *APOE* 4 carriers.

## Acknowledgements

This study was funded by NIA K01AG062683 (J.TCW.), New York Stem Cell Foundation (J.TCW.-Drunkenmiller fellowship), U01AG058635 (A.M.G), the JPB foundation (A.M.G., D.M.H.), NIA AG016573 (W.W.P.), Alzheimer’s Orange County AOC-207373 (W.W.P.), NIH NS090934 (D.M.H.), NIH AG047644 (D.M.H.), Cure Alzheimer’s Fund (F.R.M.), U01AG046170 (B.Z.), RF1AG057440 (B.Z.) and RF1AG054014 (B.Z., A.M.G.). We thank Washington University in St. Louis and University of California, Irvine Alzheimer’s disease research centers (ADRC) for providing source fibroblasts and/or hiPSCs. We thank Jill K. Gregory for drawing the schematics figure, Melanie Oaks and Seung-Ah Chung at the UCI Genomics High-Throughput Facility for their assistance with performing the RNAseq, Santiago Sole Domenech at Weill Cornell Medical College, Aurora Scrivo and Ana Maria Cuervo at Albert Einstein College of Medicine for the discussion of cell culture condition of lipid assays with respect of lysosome and autophagic function.

## Author Contributions

J.TCW., W.W.P, and A.M.G. conceived the study. J.TCW., W.W.P, and A.M.G. designed the study. J.TCW. and M.J.C. ran genetic analysis. J.TCW. performed most of the experiments and analyzed the data, assisted by W.W.P., S.B., M.J.C., M.K, E.M., and all the rest of people; J.TCW. and S.A.L. differentiated four cell types from hiPSCs. J.TCW. advised by M.K. performed genetic analysis. J.TCW. and S.B. performed global transcriptomic analysis; J.TCW. and M.W. performed human brain transcriptomic analysis; N.H.P. and J.TCW. performed the lipid assays. Y.S. from the lab of D.M.H. prepared mouse primary glia and stained by markers, and J.TCW. sequenced and analyzed the data. J.TCW., W.W.P. and A.M.G wrote the manuscript. All authors discussed the results and commented on the manuscript.

## Additional information

**Correspondence and requests for materials** should be addressed to J.TCW. or A.M.G.

