## Supplementary material for "Cholesterol and matrisome pathways dysregulated in human *APOE* ε4 glia": Methods

### Online Methods

#### Global screening array (GSA) and analysis

Genetic variant screening was performed on a GSA chip (Illumina Infinium Global Screening Array) using harvested genomic DNA (Qiagen, DNeasy Blood & Tissue Kit). To ensure high quality genotype data, samples passed the quality control (QC) metrics (SNP call rate > 95%, minor allele frequency (MAF) > 5%, Hardy-Weinberg equilibrium p-value > 1E-6, sample call rate > 95%) were analyzed. Sample ancestry was confirmed by multidimensional scaling (MDS) using PLINK 1.9[1]; MDS values for each subject were compared to 1000 Genome Project (Phase 3) sample values. Related samples were determined based on pair-wise identity-by-descent (IBD) estimation using PLINK. Duplicated samples with PI-HAT values between 0.99 to 1 were identified (see Supplementary Table 1 and 6).

#### Genetic risk score (GRS) analysis for the study line selection

GRSs were obtained from GSA and Taqman genotype based imputation results from subjects without *APOE* genotype. AD GWAS IGAP reference SNPs with their associated β[2] were computed using PLINK and the following formula:

$$\mathbf{GRS = \Sigma_j (S_j \times G_j) / M}$$

, where M is number of markers, S is β, and G is subject genotype.
We controlled the number of M = 28 from AD GWAS. The scored subjects are independent from the AD GWAS study used to generate the original weighted score.

#### Generation of hiPSC-astrocytes

The consent for reprogramming human somatic cells to hiPSC was carried out on ESCRO protocol 19-04 at Mount Sinai (J.TCW.), 2013-9561 and 2017-1061 at UCI (W.W.P.). hiPSCs maintained on Matrigel (Corning) in mTeSR1 (StemCell Technologies) supplemented with 10 ng/ml FGF2 StemBeads (StemCultures) were differentiated to NPCs by dual SMAD inhibition (0.1μM LDN193189 and 10μM SB431542) in embryoid bodies (EB) media (DMEM/F12 (Invitrogen, 10565), 1x N2 (Invitrogen, 17502-048), and 1x B27-RA (Invitrogen, 12587-010)). Rosettes were selected at 14 DIV by Rosette Selection Reagent (StemCell Technologies) and patterned to forebrain NPCs with EB media containing 20 ng/ml FGF2 (Invitrogen). NPCs ($CD271^-$/$CD133^+$) were enriched by magnetic activated cell sorting (Miltenyi Biotec)[3] and validated immunocytochemically using SOX2 (Cell signaling, 3579S), PAX6 (Abcam, ab5790), FOXP2 (Abcam, ab16046) and NESTIN (Abcam, ab22035). Dissociated single cell forebrain NPCs (15,000 cells/$cm^2$) were differentiated to astrocytes in astrocyte medium (ScienCell, 1801) on Matrigel as described[4]. Cells were continually passaged at 95% confluency and harvested as astrocytes at 30 DIV, validated immunocytochemically and/or by FACS for the astrocyte-specific markers and used for subsequent experiments.

#### Generation of hiPSC-mixed cortical cultures

Forebrain NPCs were dissociated with Accutase (Millipore) and re-plated on Matrigel at 42,000 cells/$cm^2$. After 24 h, media was replaced with BrainPhys media (StemCell Technologies) supplemented with 1% Antibiotic/Antimycotic, 1x N2, 1x B27, 20ng/ml brain-derived neurotrophic factor (BDNF), 20ng/ml glia-derived neurotrophic factor (GDNF), 250ug/ml dibutyryl cyclic AMP sodium salt (cAMP) (Sigma) and 200μM L-Ascorbic acid (AA) (Sigma) as described[5]. Cells cultured for six weeks were used for all experiments.

**Generation of hiPSC-microglia**

hiPSCs cultured in either mTeSR or E8 (StemCell Technologies) were differentiated to hematopoietic progenitor cells (HPCs) and subsequently to microglia. Single cell hiPSCs were cultured in hypoxia (20% $O_2$, 20% $CO_2$) at 37°C with 50ng/ml FGF2 (Peprotech) and 50ng/ml BMP4 (Peprotech), 12.5ng/ml Activin A (Peprotech), 2mM LiCl (Peprotech) during days 0-2 and 50ng/ml FGF2 and 50ng/ml VEGF (Peprotech) for days 2-4 in basal HPC medium (50% IMDM (Gibco), 50% F12 (Gibco), 0.02 mg/ml insulin (Sigma), 2% v/v ITSG-X (Gibco), 64μg/ml ascorbic acid (Sigma), 400 μM monothioglycerol (MTG) (Sigma), 10 μg/ml PVA (Sigma), 1x GlutaMax (Gibco), 1x Chemically-defined lipid concentrate (Gibco), 1x non-essential amino acids (NEAA) (Gibco), 1% v/v Antibiotic/Antimycotic) to generate EBs as described previously[6]. EBs were transferred to normoxia for another six days in basal HPC media supplemented with 50ng/ml of each FGF2, VEGF, TPO and IL6, and 10ng/ml of each SCF and IL3. At day 10, HPCs were collected in the absence of FACS, filtered through a 45um cell strainer (Fisher Scientific), and plated onto Matrigel-coated 6-well plates (200,000 cells/well). HPCs were cultured in microglia basal medium (DMEM/F12, 1x Glutamax, 1x NEAA, 2% v/v ITS-G, 2% v/v B27, 0.5% v/v N2, 200uM MTG, 5ug/ml Insulin) with 50 ng/mL TGFβ (Peprotech), 100 ng/mL IL-34 (Peprotech) and 25 ng/mL M-CSF (Peprotech) for 25 days, and then cultured in microglia basal medium supplemented with 100 ng/mL CX3CL1 (Peprotech) and 100 ng/mL CD200 (Novaprotein) for an additional three days, and harvested on day 28 for analysis. HPCs were validated by FACS for $CD43^{+}$(> 98%) and $CD41^{+}/CD235a^{+}/CD45^{-}$ (Biolegend) while microglia were validated by immunocytochemistry with microglia-specific markers.

**Generation of hiPSCs-BMEC**

BMECs were generated as described[7]. Briefly, hiPSCs were plated as evenly dispersed single cells on Matrigel (Corning)-coated 6-well plates. Cells were maintained in E8 at 37ºC in 5% $CO_2$ with daily media change until 70% confluent and subsequently switched to unconditioned media (Dulbecco's Modified Eagle's Medium/Ham's F12 (Gibco) with 20% Knock-Out Serum Replacement (ThermoFisher), 1% NEAA, 0.836 μM beta-mercaptoethanol (MilliporeSigma) and 5% GlutaMAX and cultured until a monolayer exhibiting distinct morphological changes formed (3-5 days). Next, cells were cultured in basal EC media (Human Endothelial Serum-Free Media (ThermoFisher) with 1% human platelet poor derived serum (MilliporeSigma)) supplemented with 20 ng/mL bFGF (Peprotech) and 10 μg/mL retinoic acid (RA) (Sigma) for 24 h, followed by incubation in EC media lacking bFGF for another 24 h, and then cultured in basal EC media for an additional 48 h. Afterward, cells were dissociated into single cells (Tryple Express (Gibco)) and plated onto either 24-well plates coated with a collagen IV (400 μg/mL; Sigma)/fibronectin (100 μg/mL; Sigma) in $H_2O$ mixture overnight in an incubator. Resultant BMECs were cultured in EC media for an additional 24 h prior to experiments.

**Karyotyping**

Karyotyping was performed by Wicell Cytogenetics (Madison, WI).

**Purification of primary astrocytes and microglia**

All animal procedures and experiments were performed under guidelines approved by the animal studies committee at Washington University School of Medicine (D.M.H. protocol: 20180139). Mixed glia were obtained at P2 from human *APOE* targeted replacement mice (C57BL/6, provided by Dr. Patrick M. Sullivan) or *Apoe* knockout mice (C57BL/6, Jackson Laboratory, 002052). Mouse cortices were dissected in Hanks' Balanced Salt solution (HBSS without $Ca^{2+}$, $Mg^{2+}$). After meninges removal, tissue was digested in 0.25% trypsin (GIBCO, 15090-046) and 0.2mg/ml DNase (Sigma, DN-25) at 37 °C for 10 min, washed with HBSS, and dissociated in HBSS containing 0.4mg/ml DNase using fire-polished Pasteur glass pipettes followed by filtration through a 70-µm nylon mesh. Cells were pelleted (1,000 g, 5 min), washed with glial medium (DMEM, 10% FBS, 1x Pen/strep, 1x Glutamax), and plated onto 10 µg/ml Poly-L-lysine (Sigma, P2636)-coated 10cm tissue culture dishes in glial medium, followed by medium replacement on the second day. After cells reached confluency, they were cultured for an additional 5-7 days with fresh glial media supplement to allow for microglia proliferation atop astrocytes. Microglia were flushed off the astrocyte layer via pipetting and plated (62,500 cells/cm$^2$) in glial medium. Astrocytes were collected by trypsinization and replated (52,000 cells/cm$^2$). Astrocytes and microglia were cultured in hiPSC-astrocyte (ScienCell) and hiPSC-microglia media (IL34, TGFβ, M-CSF for 4 days, followed by CX3CL and CD200 for 3 days) for 7 days respectively to mimic the human cell culture conditions.

**Immunohistochemistry**

Cells were fixed in 4% paraformaldehyde in PBS at 4°C for 10 min. hiPSCs and differentiated CNS cells were permeabilized (1.0% Triton in PBS) at room temperature for 15 min and blocked in 5% donkey serum with 0.1% Triton at room temperature for 30 min. The primary antibodies used for hiPSC-astrocytes were 1:1,000 anti-S100β (Sigma-Aldrich, S2532), 1:500 anti-Vimentin (Cell Signaling, R28#3932), 1:500 anti-NFIA (Active Motif, 39398), 1:100 anti-GLAST/EAAT1 (BOSTER: PA2185), 1:100 anti-ALDH1L2 (Novusbio, NBP1-81935) and 1:500 anti-AQP4 (Almone Labs, AQP-004), for hiPSC-mixed cortical cultures were 1: 400 anti-MAP2AB (Sigma, M1406), 1: 1,000 TUJ1 (Biolegend, 802001), 1:500 anti-TH1 (Pel-Freez Biologicals, P40101) and 1:500 anti-GABA (Sigma; A2052), for hiPSC-microglia were 2µg/ml anti-CX3CR1 (BioRad, AHP1589), for both hiPSC-microglia and mouse microglia were 1:300 anti-IBA1 (Sigma, MABN92), 10ug/ml anti-TREM2 (R&D, AF1828), 1:1,000, anti-P2RY12 (Sigma, HPA014518) and 1:100 PU.1 (Cell Signaling, 2266) and for hiPSC-BMECs were 1:100 anti-Claudin-5 (ThermoFisher, 4C3C2), 1:200 anti-ZO-1 (ThermoFisher, 402200) and 1:200 anti-Occludin (ThermoFisher, OC-3F10). For mouse astrocytes, 1:1,000 anti-GFAP (Millipore, MAB3402), 1:400 anti-EAAT1 (GeneTex, GTX134060) and 1:1,000 anti-ALDH1L1 (Abcam, ab190298). For hiPSC, anti-Nanog (Cell signaling, 4903S), anti-OCT4 (Cell signaling, 2840S), anti-TRA1-60 (Cell signaling, 4746P) and anti-TRA1-81 (Cell signaling, 4745P). Secondary antibodies used were 1:300 Alexa donkey 488 and 568 anti-rabbit, mouse, or chicken (Life Technologies). DAPI (4',6-diamidino-2-phenylindole, 0.5 µg/ml) was used to visualize nuclei. Images were acquired using an Olympus IX51 Fluorescence Microscope, a Zeiss LSM780 confocal microscope or Cytation 5 imager.

**RNAseq analysis of hiPSC-derived brain cells and primary mouse glia**

Cells were harvested, stored in RNAlater, and RNA subsequently extracted from all cell types using RNeasy Mini (Qiagen) following manufacturer's guidelines. Deep RNA sequencing (50-70 million reads per sample) was performed by UCI Genomics High-Throughput Facility for hiPSC-brain cells and by GeneWiz for mouse primary glia. The Illumina TruSeq mRNA stranded protocol was used to isolate poly-A mRNA from RNA (RNA integrity score $\geq 9$) and 200 ng was used to construct libraries that were quantified and normalized using the Library Quantification Kit (Kapa

Biosystems) and sequenced as paired-end 100 bp reads on the Illumina HiSeq 4000. Paired-end sequencing reads were aligned to the hg38 genome using Star aligner[8]. Aligned and sorted bam files were loaded into IGV to verify *APOE* genotype[9]. FeatureCounts[10] was used to quantify gene expression based on GENCODE. Gene level read counts were normalized as Counts per Million mapped reads using trimmed mean of M-values (TMM) normalization[11] to adjust for sequencing library variance. The ERCC spike-in control[12] was used to adjust for sequencing batch effect. Sex effects were corrected by linear regression. Multi-dimensional scaling and cluster analysis was performed using R. Next, DEGs between different *APOE* genotypes within the same cell types were identified by linear model analysis using the Bioconductor package DESeq2[13]. For multiple test adjustment, the false discovery rate (FDR) of the differential expression test was estimated using the Benjamini–Hochberg method[14]. To test if known gene ontology (GO) and pathways are enriched for DEGs, fast preranked gene set enrichment analysis (fGSEA) was performed using GO annotations and canonical pathways (Biocarta, KEGG and Reactome) available from the Molecular Signatures Database (MSigDB)[15], following 1 million gene set permutations. For mouse data, homology conversion of mouse to human DEGs are carried out, followed by fGSEA. The functional and causal network analyses were further performed by integrating DEGs of significantly enriched pathways into Ingenuity Pathway Analysis (IPA)[16]. Significantly enriched pathways and disease/functional annotations were identified (Supplementary Table 2).

**RNAseq analysis on human postmortem brain and deconvolution analysis**

DEG analysis on postmortem brain was stratified by region and disease severity: prefrontal cortex (PFC, BA10), superior temporal gyrus (STG, BA22), parahippocampal gyrus (PHG, BA36) and inferior frontal gyrus (IFG, BA44)[17] and CDR (0-5), clinical phenotype (Cerebral AD, possible AD, probable AD and definite AD) and neuropathological plaque severity (normal, mild, medium and severe) and further plotted with significant pathways of FDR $< 0.05$. For deconvolution analysis, digital sorting algorithm (DSA)[18], population-specific expression analysis (PSEA)[19], and non-negative matrix factorization (ssKL)[20] and a PCA-based method modified from CellCODE (BRETIGEA, BRain cEll Type specIfic Gene Expression Analysis)[21] were tested on available RNAseq data from human primary brain cells[22], hiPSC-derived cells[4] and our dataset. Given the relatively clear cell type composition of the public data, they served as a reference to determine the best method and parameters. Each method required known brain cell type-specific markers to either infer the cell type composition or compute the composition-informative surrogate variables. We used a consensus top ranked cell type-specific marker set from a recent meta-analysis of multiple cell type-specific and single cell RNAseq datasets[21]. Performance was evaluated using a series of marker numbers (1, 3, 5, 10, up to 50). DSA (marker size 5 and 10) performed best and therefore was used in all subsequent deconvolution analyses. We used brain RNAseq data from the Mount Sinai Brain Bank AD[23] and ROSMAP[24] cohorts (Supplementary Table 4). Before cell type proportion inference, we corrected for: postmortem interval (PMI), age of death (AOD), RIN, exonic rate, race, rRNA rate, sex and batch. We deconvoluted the data for each brain region separately using csSAM[25] and subsequently performed DEG analysis between *APOE* 44 and *APOE* 33 in each cell type stratified by disease status measured by CDR (0, 0.5, 1, 2, 3, 4 and 5). GO and pathways enriched for DEGs were computed by fGSEA[15]. Significantly enriched pathways were further investigated by IPA[16]. Significantly enriched pathways and disease/functional annotations were identified (Supplementary Table 3).

**Meta-analysis of brain cell type specific RNAseq data (Zhang et al. 2016)**

Downloaded raw RNAseq data of multiple brain cell types from gene expression omnibus (accession GSE73721)[22] was processed using the same star-FeatureCounts pipeline described above. The gene level read counts were merged with the data generated in this project and normalized using the TMM approach and sex and batch corrected using linear regression. Hierarchical cluster analysis was performed using R.

**Generation of isogenic CRISPR/Cas9 gene-edited hiPSCs.** TCW1 and TCW2 (Supplementary Table 6) were generated as described[26] by the 'CORRECT' scarless gene-editing method[27]. Briefly, the CRISPR Design tool (http://crispr.mit.edu) was used to identify the sgRNA 5'– CCTCGCCGCGGTACTGCACCAGG – 3', which was cloned into the pX330-U6-Chimeric_BB-CBh-hSpCas9-GFP (PX338) plasmid (Addgene, 42230). The correct *APOE* sgRNA sequence orientation was confirmed by Sanger sequencing and CRISPR/Cas9-*APOE* sgRNA plasmid cleavage efficiency was determined using the Surveyor mutation detection kit in 293T cells. The single-strand oligo-deoxynucleotide (ssODN) was designed to convert *APOE* ε4 to *APOE* ε3 with a protospacer adjacent motif (PAM) silent mutation to prevent recurrent Cas9 editing. hiPSCs (70-80% confluent) dissociated by Accutase supplemented with 10 uM Thiazovivin (Tzv) (Millipore), were harvested (200 x g, 3 min), and electroporated (Neon®, ThermoFisher) according to the manufacturer's instructions. In brief, cells resuspended in 10µl Neon Resuspension Buffer R, 1µg CRISPR/Cas9-*APOE* sgRNA plasmid and 1µl of 10µM of ssODN were electroporated plated on Matrigel-coated plates in mTeSR media with 10 µM Tzv for 72h. GFP-expressing hiPSC were isolated by FACS (BD FACSAria). Sorted single cells were suspended in mTeSR with Tzv and plated into 96 well plates containing MEFs (4,000 cells/well). Clones were expanded and transferred to a replicate plate for gDNA isolation and Sanger sequencing to identify genome edited clones.

**Cellular cholesterol determination by Gas Chromatography-Mass Spectrometry (GC-MS)**
Isogenic *APOE* astrocytes were plated (200,000 cells/well of 6-well-plate) in serum-free astrocyte media. After 48 h, cells were washed twice with HBSS, and cellular lipids extracted with hexane-isopropyl alcohol (3:2, v/v)[28] and dried under Argon stream. Lipids were resuspended in hexane and analyzed by GC-MS on a HP 5890 series II gas chromatograph (Hewlett-Packard) equipped with a flame-ionization detector. Lipids were separated on a HP-5 capillary column (15 m × 0.53 mm) coated with 5% phenyl methyl siloxane (1.5 µm) in which the injection temperature was maintained at 255°C, the oven temperature was isothermally held at 260°C and using a helium mobile phase (30 ml/min flow rate). The astrocyte unesterified/or free cholesterol (FC) content in each well was quantified using β-sitosterol as an internal standard. After lipid extraction, cells were lysed in 0.1M NaOH, and the protein content of each well (µg protein/well) was determined using the DC protein kit (Bio-Rad). The FC content of each well was normalized to the corresponding protein concentration. Experimental data are presented as fractions of control.

**Intracellular FC measurement by filipin stain and image analysis**
After isogenic *APOE* astrocytes were plated (20,000 cells/ $cm^2$) in serum-free astrocyte media, cells were treated with or without LDL for 24 h, and intracellular FC was determined by fluorescence. Cultured astrocytes were fixed, and FC were labeled with filipin (50 µg/ml in PBS for 45 min at room temperature), followed by 3x wash with PBS, and acquiring images (20 images per experiment, 10-20 cells per image) by widefield fluorescence microscope (Leica Microsystems, Germany) using a 20x objective and standard A4 UV filter with 17% neutral density to measure

filipin fluorescence intensity. In each experiment 20 images were acquired, and each image had 10-20 cells in the field. In order to avoid bias, different pairs of *APOE* 33 and *APOE* 44 clones were used. Thus with three clones in each of the *APOE* 33 and *APOE* 44 lines, nine combinations were tested. Experiments were repeated at least twice per pair. For imaging, cellular FC content was determined using Metamorph Discovery-1 image-analysis software. Background was subtracted from each shading-corrected image by determining the fifth percentile intensity value of the image and subtracting this value from each pixel in the image. At the plating density used, all fields were 50-70% confluent in imaged areas. Next, a low threshold was applied to include all areas occupied by cells. The outline of cells using the selected values were comparable to cell outlines in transmitted light images. For the average cellular FC, the low threshold was used to measure total filipin intensity above the threshold and was divided by the number of pixels above the lower threshold for each field to yield the average filipin intensity/cell area. All data were normalized to *APOE* 33 control within each experiment.

**Lysosomal Localization**

After 24 h incubation of isogenic astrocytes (20,000 cells/$cm^2$) in serum-free astrocyte media, cells were labeled with 2 mg/ml FITC-Dextran and incubated for an additional 24 h to allow endocytosis and delivery to lysosomes. Subsequently, FITC-Dextran (70 KDa) labeled astrocytes were chased with fresh astrocyte media for 2 h, washed with PBS, fixed with 2% PFA, and stained with 50 µg/ml filipin as described above. Images were acquired on Zeiss LSM 880, AxioObserver confocal microscope equipped with a Plan-Apochromat 63x oil DIC M27 objective and UV 405 and HeNe 543 lasers. Z-stacks were acquired using a step size equivalent to one airy unit. Six images with sum projections were acquired per sample.

**Western blot**

Protein lysates were collected at 4 °C in RIPA buffer (Sigma) supplemented with protease inhibitors (10uM leupeptin, 5ug/ml pepstatin A, 3ug/ml aprotinin, 25ug/ml ALLN, and 0.5mM PMSF). Conditioned growth medium was collected at 4 °C with protease/phosphatase inhibitors (Cell Signaling) and concentrated with Amicon Ultra-15 filters (30-kDa cut-off, EMD Millipore). Total protein concentration was determined by BCA method (Thermofisher Scientific) and equal amounts of total protein were loaded onto 8% or 4-12% Bis-Tris Plus Gels (Thermofisher Scientific). Following electrophoresis (100 V, 1h), proteins were transferred to iBlot® 2 Transfer Stacks, nitrocellulose membranes (Thermofisher Scientific). Blots were probed overnight at 4 °C with 1:500 anti-HMG-CoA reductase (EMD Milipore, ABS229), 1:1,000 anti-APOE (Calbiocham, 178479), 1:1,000 anti-LAMP1 (Abcam, ab24170), or 1:700 anti-ABCA1 (Abcam, ab18180) followed by 1:2,000 HRP-conjugated secondary (Goat, life technologies, 611620; Rabbit, Vector Laboratories, PI-1000; Mouse, Vector Laboratories, PI-2000, 1 h at room temperature) and visualized with WesternBright™ ECL HRP Substrate reagents (Advansta) on the UVP System.

**Multiplex immunoassay and analysis**

Isogenic *APOE* astrocytes incubated in serum free-conditioned media (20 h) were harvested with protease/phosphatase inhibitors, centrifuged (400 g, 4 min), and supernatant collected. Cell number was determined in order to load equal total protein. Samples were loaded in 96 well plates with standard controls containing cytokine/chemokine/growth factor 45-Plex Human Panel 1 (Invitrogen, ProcartaPlex) following manufacturer guidelines. Each well was loaded and quantified for each protein using the Luminex system (Magpix and xPONENT4.2)[29]. Hierarchical clustering and correlation coefficient analyses of differentially secreted proteins were performed using R.

### Statistical analysis

Statistical analyses were performed using Prism 5 (GraphPad Software). One-way ANOVA followed by Bonferroni's Multiple comparison post hoc test or unpaired one-tailed Student's t tests were used. Data are represented as mean ± SEM.

### Data availability

The data that support the findings of this study will be available from dbGaP once the manuscript is accepted for publication.

### Code availability

Although we have used the software cited in this manuscript with default parameters or minor changes, code for these analyses is available upon request.
