## Extended Data Figures 1-6 for "Cholesterol and matrisome pathways dysregulated in human *APOE* ε4 glia"

**Extended Data Figure 1. GWAS and *APOE* genotype based subject selection and four brain cell type differentiation**

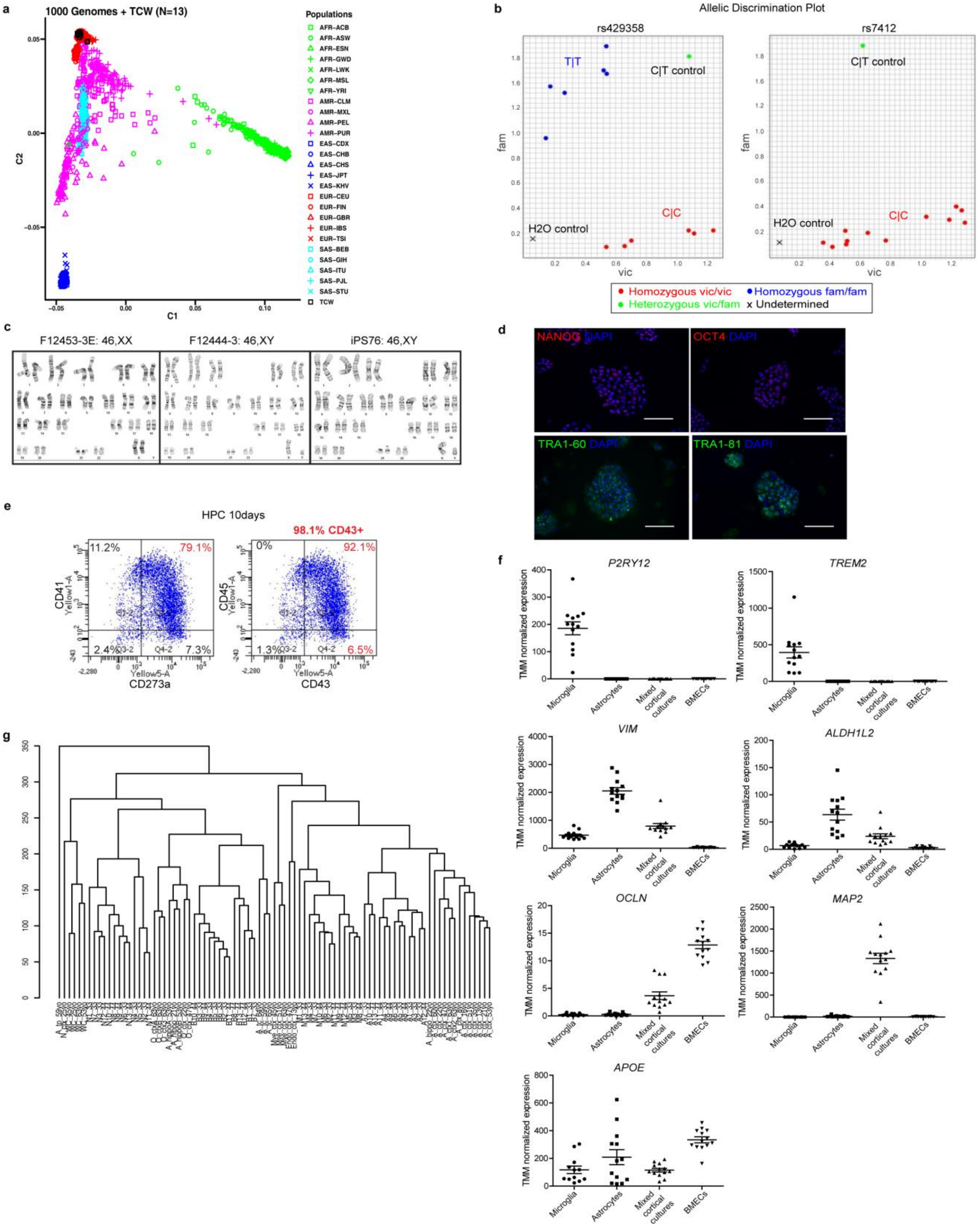

**a.** Ethnic background of lines used in this study (labeled TCW, black) from GSA compared to the 1000 Genomes reference confirmed all individuals were of European descent. **b.** Allelic discrimination plot of *APOE* SNPs, rs429358 and rs7412, of the 13 individual lines. **c.** Representative normal karyotype results of hiPSC lines. **d.** Validation of pluripotency in hiPSC lines: representative immunofluorescence images confirming expression of the pluripotency markers NANOG, OCT4, TRA1-60 and TRA-1-81 from hiPSC lines derived from fibroblasts. **e.** FACS plots of HPCs at 10-day of differentiation. The selected population used for microglia differentiation are >98% CD43+ with 86.4% CD273a+, 90.3% CD41+ and 92.1% CD45+. **f.** Relative expression levels of cell type specific markers from TMM normalized counts of RNAseq data in the four CNS cell types. **g.** Cluster analysis of hiPSC-derived cells compared to primary human brain cells from Zhang et al. 2016. A, astrocytes; B, BMECs; M, microglia; N, mixed cortical cultures; WC, whole cortex; O, oligodendrocytes; Mye, myeloid cells; Endo, endothelial cells; tp, temporal lobe; hippo, hippocampus; ctx, cortex.

**Extended Data Figure 2. Significant cell autonomous and non-cell autonomous effects of *APOE 44* in hiPSC-microglia, astrocytes and mixed cortical cultures.**

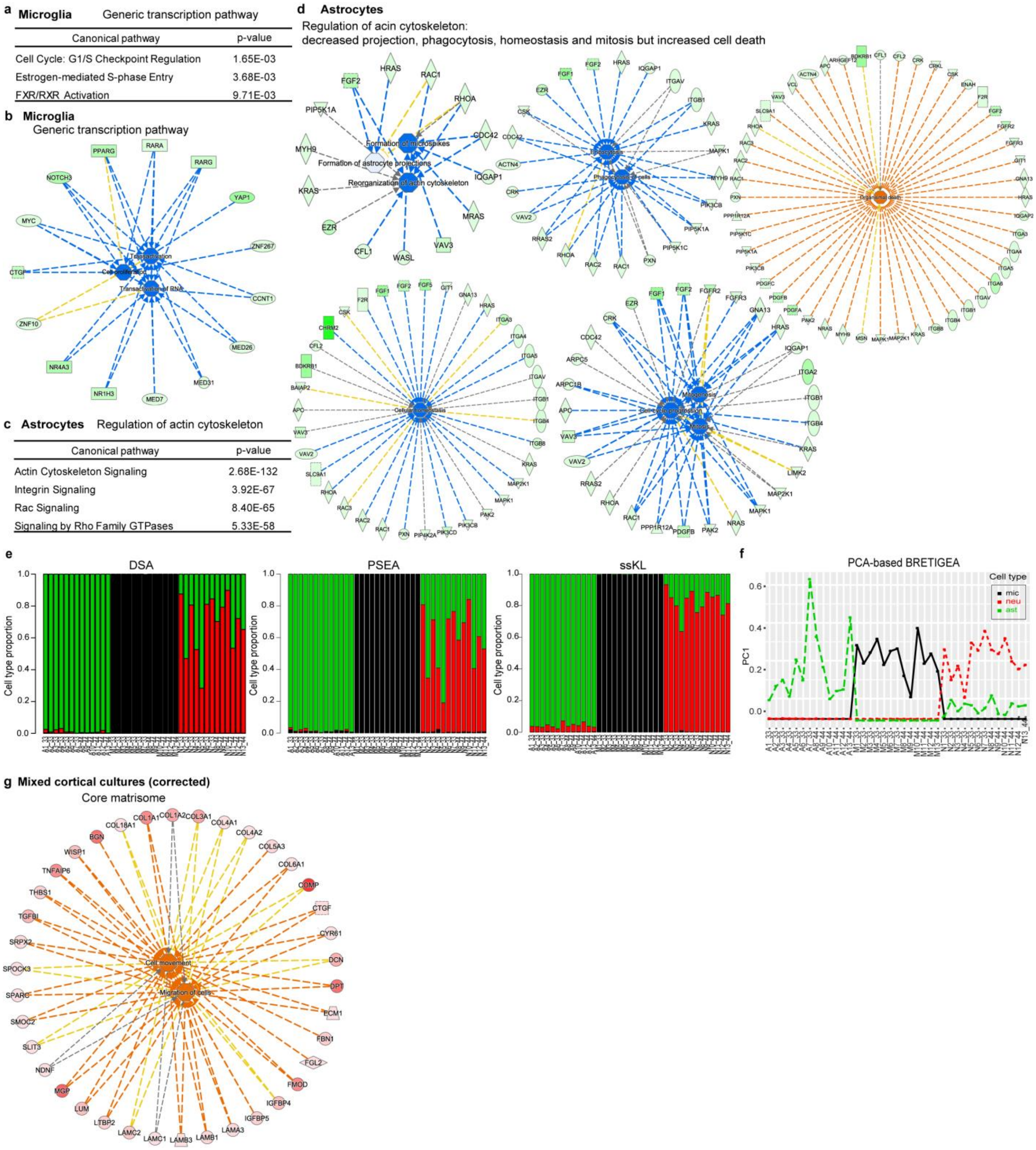

**a.** Top canonical pathway analysis of Generic transcription pathway, negatively enriched in *APOE* 44 microglia from GSEA. **b.** Molecular functions of Generic transcription pathway, enriched in *APOE* 44 microglia. **c.** Top canonical pathway analysis of Regulation of actin cytoskeleton, negatively enriched in *APOE* 44 microglia from GSEA. **d.** Molecular functions of Regulation of actin cytoskeleton, enriched in *APOE* 44 astrocytes. **e-f.** Cell type proportion analysis of hiPSC-astrocytes, microglia and neurons by DSA, PSEA and ssKL (**e**) and by a PCA-based method modified from CellCODE (BRETIGEA) (**f**). **g.** Functional pathway analysis by ingenuity pathway of Core matrisome, enriched in *APOE* 44 compared to *APOE* 33 in cell type proportion corrected hiPSC-mixed cortical cultures.

**Extended Data Figure 3. Brain cell type deconvolution and comparison of *APOE* 44 vs *APOE* 33 and disease phenotypes vs control.**

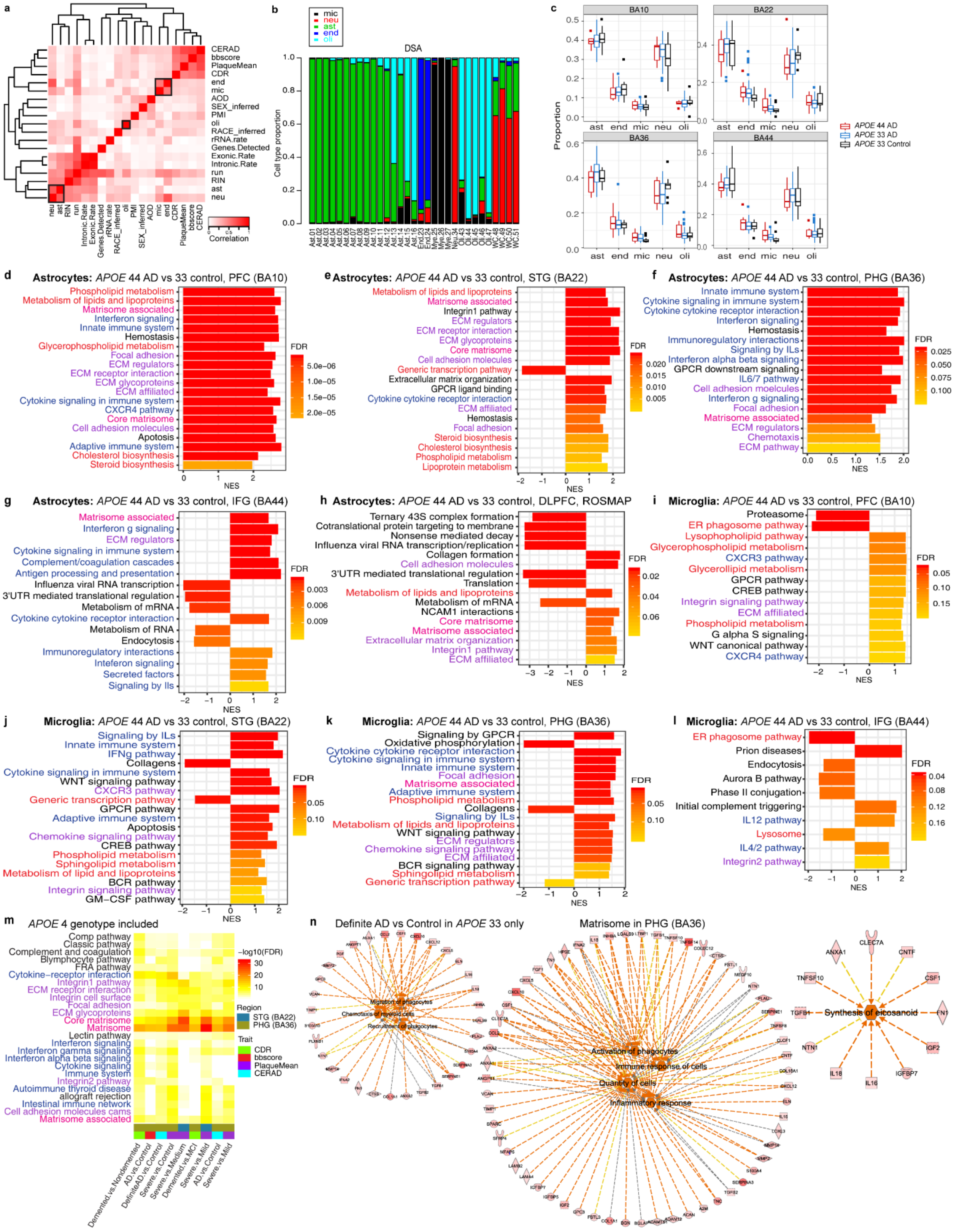

**a.** Spearman correlation and cluster analysis of all confounding factors including five cell types (marked as black boxes). **b.** Cell type deconvolution of primary human cell types, neurons, astrocytes, microglia, endothelial cells and oligodendrocytes using DSA algorithm. **c.** Cell type proportion changes by *APOE* genotype in AD and control brains. Each comparison of t-test statistics (p-values) are listed in **Supplementary table 5**. **d-l.** GSEA of DEGs of *APOE* 44 compared to *APOE* 33 in each cell type (astrocytes or microglia) after cell type deconvolution in various regions of AD brain from multiple cohorts (MSBB and ROSMAP). **m.** Upregulated canonical pathways of DEGs in different regions of brain comparing various criteria of AD-related phenotypes including *APOE* 4 carriers. Functional pathway statistics of Matrisome associated, Matrisome and Core matrisome (marked with black boxes) were analyzed in **Supplementary Table 3**. **n.** Functional pathway analysis by ingenuity pathway of Matrisome, enriched in definite AD vs control only in *APOE* 33 groups.

**Extended Data Figure 4. Lipid metabolic dysfunction is specific to human *APOE* 44 glia, which partially resemble mouse *Apoe* knockout but not *APOE* 44 glia**

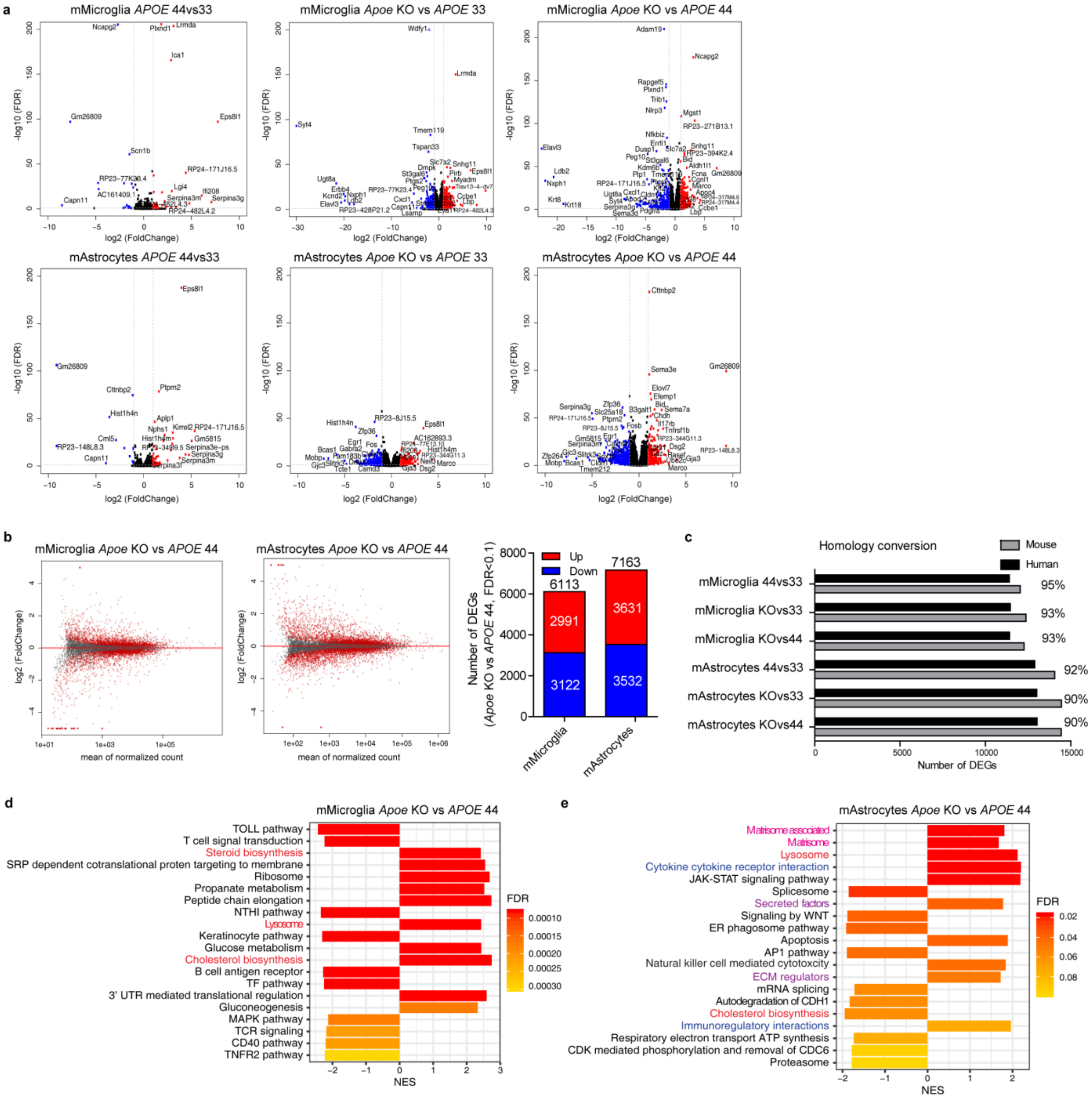


**a.** Volcano plots comparing mouse microglia (mMicroglia) or astrocytes (mAstrocytes) of different *APOE* genotypes (targeted replacement mice: *hAPOE* 44, *hAPOE* 33 and *mApoe-/-*). Average log2 (fold change) by log10 (FDR) is shown for all genes. Genes upregulated (red) and downregulated (blue) by > |2.5| - |4| fold change with FDR <0.1 are labeled with gene names. **b.** MA plots of *Apoe* KO vs *APOE* 44 in mMicroglia and mAstrocytes (left) and the number of significant DEGs in MA plots (right). **c.** Homology conversion ratio of mouse to human DEGs in each different *APOE* genotype comparison in mMicroglia and mAstrocytes. **d-e.** Gene set enrichment analysis of DEGs of *Apoe* KO compared to *APOE* 44 in mMicroglia (**e**) and mAstrocytes (**f**).

**Extended Data Figure 5. Generation of isogenic *APOE* hiPSC lines and free cholesterol level in fibroblasts**

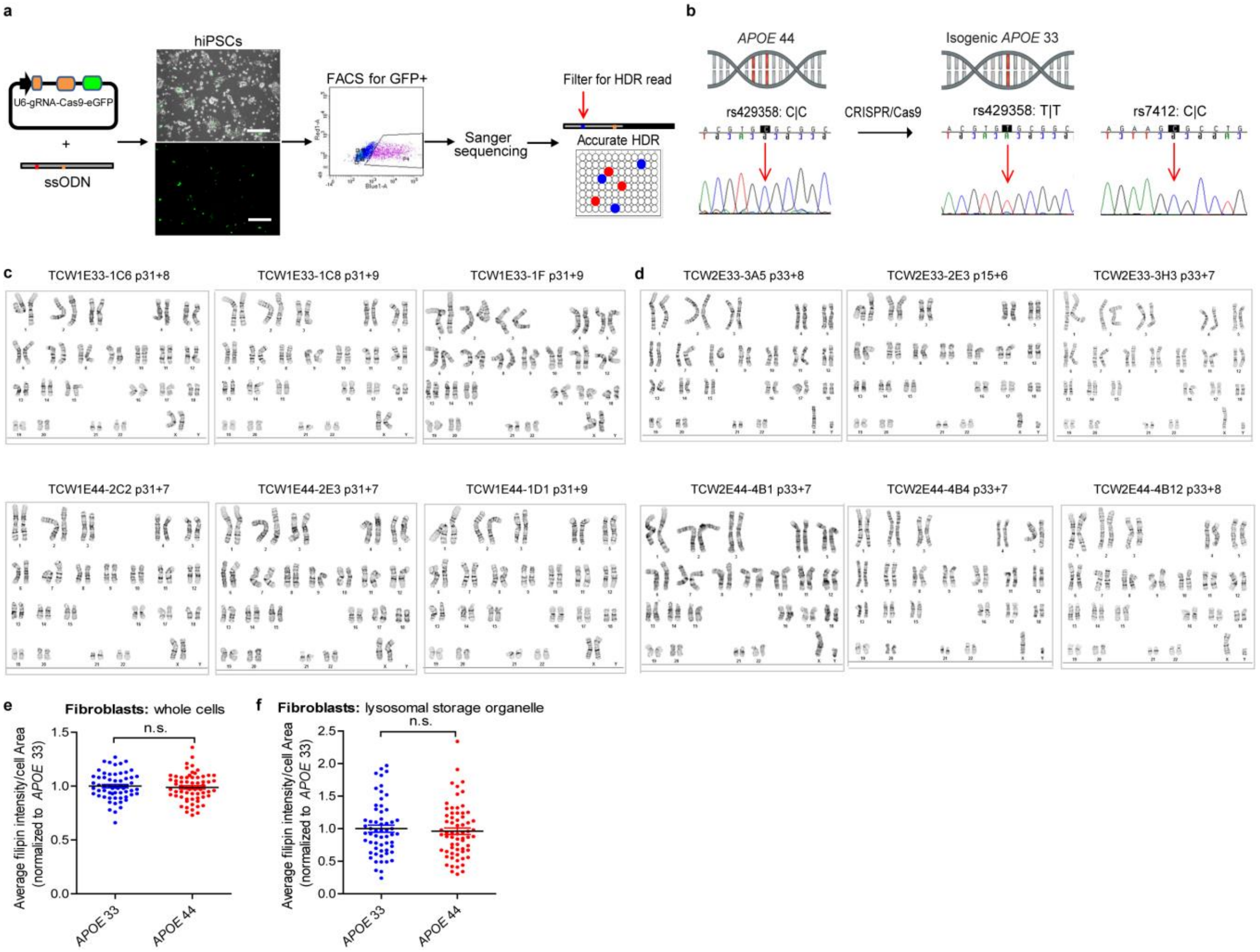

**a.** Schematics of generation of CRISPR/Cas9 genome edited *APOE* hiPSC lines. HDR, homology DNA repair. **b.** DNA sequence of the *APOE* locus at rs429358 and rs7412 in genomic DNA from hiPSCs, confirming the homozygous conversion from C|C (left) to T|T (right) at rs429358, which determines *APOE* 44 and *APOE* 33**. c-d.** G-banding karyotype results of each genome edited hiPSC line (TCW1 from individual ID 8 and TCW2 lines from individual ID 10) after *APOE* genotype confirmation. Identities of each lines with the original source fibroblasts were confirmed by GSA in **Supplementary Table 6**. **e-f.** Average filipin level in *APOE* 33 and *APOE* 44 fibroblasts of whole cells (e) and lysosomal storage organelle (f).

**Extended Data Figure 6. Secreted proteins from isogenic *APOE* 44 astrocytes show activated profiles of chemokines and cytokines compared to *APOE* 33 cells.**

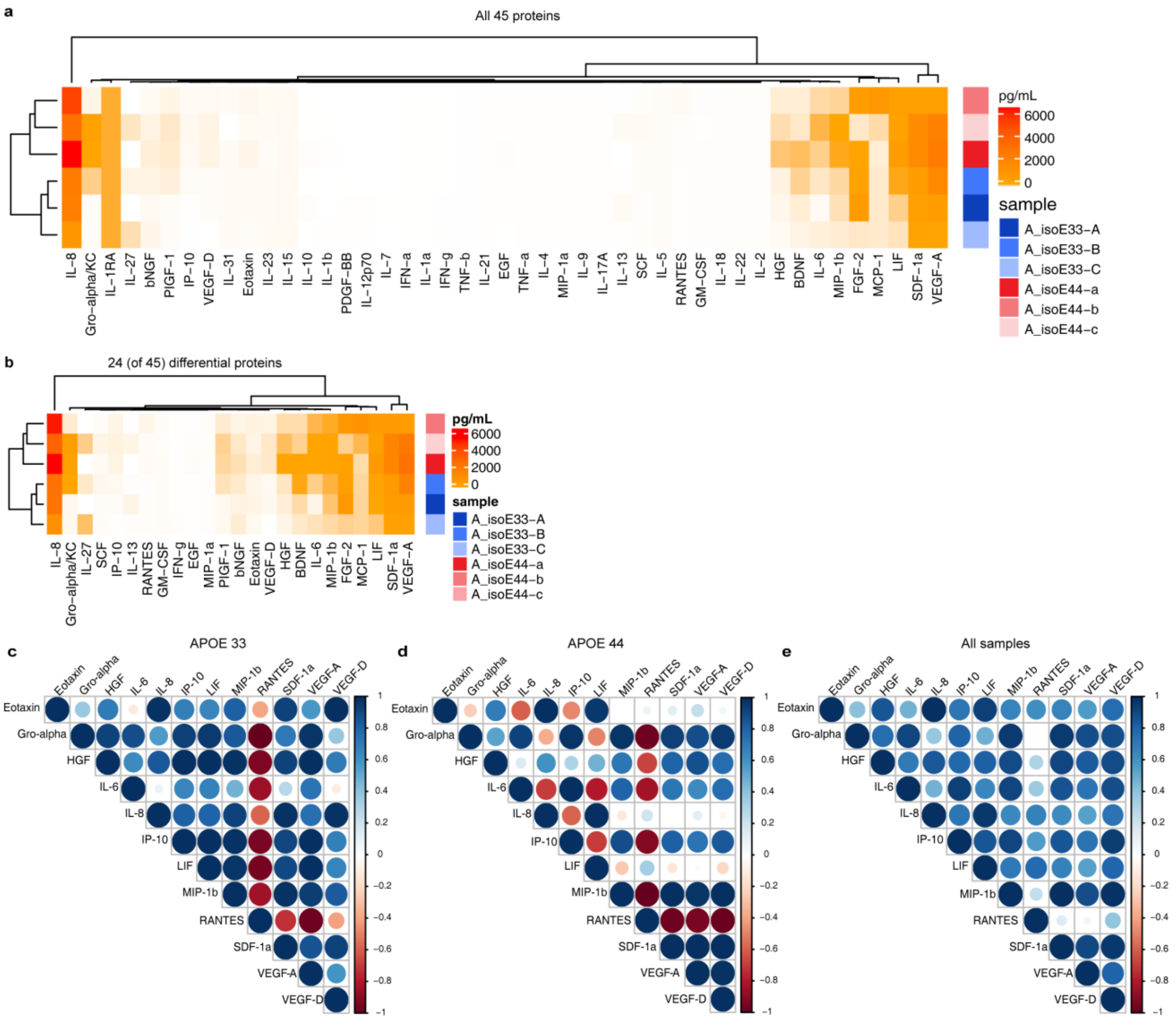


**a.** Clustering heatmap for the 45-plex human panel 1 including chemokines, cytokines and growth factors. **b.** Clustering heatmap for the top 24 differentially secreted proteins by *APOE* genotype. A_isoE33, isogenic *APOE* 33 astrocytes, A_isoE44, isogenic *APOE* 44 astrocytes. A-C and a-c are independent genome edited CRISPR lines. **c-e.** Correlation coefficient analysis for the top 12 differentially secreted proteins in *APOE* 33 (**c**), *APOE* 44 (**d**) and all samples (**e**).
