## Supplementary Information for "Cholesterol and matrisome pathways dysregulated in human *APOE* ε4 glia"

### Supplemental Data Table 1. Genetic and clinical information of the hiPSC lines.

| Individual ID | *APOE* genotype | GRS – APOE score | Ethnicity | Age of onset | Age at skin biopsy/ PBMC collection | Gender | Disease status (CDR at biopsy) | PI HAT of fibroblast and hiPSC (GSA) |
|---|---|---|---|---|---|---|---|---|
| 1 | 33 | 0.0191 | Caucasian | - | 76 | Female | Control (0) | 0.9999 |
| 2 | 33 | -0.00046 | Caucasian | - | 67 | Male | Control (0) | 1 |
| 3 | 33 | -0.00112 | Caucasian | - | 81 | Female | Control (0) | 1 |
| 4 | 33 | 0.00772 | Caucasian | 78 | 81 | Female | AD (2) | 0.9998 |
| 5 | 33 | 0.00722 | Caucasian | 82 | 94 | Female | AD (0.5) | 1 |
| 6 | 33 | 0.01647 | Caucasian | - | 87 | Male | Control (0) | 0.9997 |
| 7 | 33 | 0.0097 | Caucasian | - | 83 | Male | Control (0) | 1 |
| 8 | 44 | 0.00343 | Caucasian | 64 | 72 | Female | AD (2) | 1 |
| 9 | 44 | 0.01086 | Caucasian | - | 76 | Female | Control (0) | 1 |
| 10 | 44 | 0.00468 | Caucasian | 77 | 80 | Male | AD (0.5) | 1 |
| 11 | 44 | 0.00772 | Caucasian | 83 | 80 | Male | AD (0.5) | 1 |
| 12 | 44 | 0.01203 | Caucasian | 61 | 72 | Female | AD (2) | 0.9999 |
| 13 | 44 | 0.00652 | Caucasian | 71 | 78 | Male | AD (3) | 0.9997 |

Abbreviations: *APOE*, apolipoprotein E; GRS, genetic risk score; AD, Alzheimer's disease; PBMC, peripheral blood mononuclear cell; CDR, clinical dementia rating; GSA, global screening array.

**Supplemental Data Table 2. Ingenuity pathway z-score of enriched pathways from DEG analysis comparing *APOE* genotype in hiPSC-microglia and astrocytes.**

**Microglia: *APOE* 44 vs 33**

| Enriched pathways | Diseases or Functions annotation | p-Value | Predicted Activation State | Activation z-score |
|---|---|---|---|---|
| Cholesterol biosynthesis | Synthesis of cholesterol | 8.63E-42 | Increased | 2.216 |
| | Metabolism of cholesterol | 1.74E-26 | Increased | 2.216 |
| | Steroid metabolism | 3.91E-23 | Increased | 2.216 |
| | Synthesis of terpenoid | 4.54E-18 | Increased | 2.201 |
| Lysosome | Accumulation of lipid | 4.9E-17 | Increased | 2.805 |
| | Accumulation of cholesterol | 9.53E-10 | Increased | 2.765 |
| | Catabolism of lipid | 4.17E-16 | Decreased | -2.764 |
| | Cleavage of lipid | 1.9E-09 | Decreased | -3.364 |
| HDL-mediated lipid transport | Efflux of cholesterol | 1.87E-14 | Decreased | -2.382 |
| | Cholesterol transport | 2.11E-16 | Decreased | -2.163 |
| | Removal of lipid | 2.79E-14 | Decreased | -2.109 |

Abbreviations: *APOE*, apolipoprotein E; HDL, high density lipoprotein

**Astrocytes: *APOE* 44 vs 33**

| Enriched pathways | Diseases or Functions annotation | p-Value | Predicted Activation State | Activation z-score |
|---|---|---|---|---|
| Cholesterol biosynthesis | Synthesis of cholesterol | 7.03E-37 | Increased | 2.216 |
| | Metabolism of cholesterol | 4.92E-20 | Increased | 2.216 |
| | Steroid metabolism | 6.43E-18 | Increased | 2.216 |
| | Synthesis of terpenoid | 3.02E-15 | Increased | 2.201 |

**Supplemental Data Table 3. Ingenuity pathway z-score of enriched pathways from the DEG analysis comparing *APOE* 44 vs *APOE* 33 in hiPSC-mixed cortical cultures, definite AD vs control in *APOE* 33 brains and severe vs mild dementia in AD brains including *APOE* 4 carriers.**

### hiPSC-mixed cortical cultures (corrected): *APOE* 44 vs 33

| Enriched pathways | Diseases or Functions annotation | p-Value | Predicted Activation State | Activation z-score |
|---|---|---|---|---|
| Matrisome associated | Chemotaxis | 2.9E-31 | Increased | 5.729 |
| | Chemotaxis of myeloid cells | 1.45E-12 | Increased | 3.976 |
| | Chemotaxis of phagocytes | 1.88E-11 | Increased | 3.721 |
| | Activation of phagocytes | 1.41E-17 | Increased | 3.513 |
| | Inflammatory response | 5.39E-26 | Increased | 4.657 |
| | Synthesis of lipid | 4.38E-16 | Increased | 2.879 |
| | Synthesis of eicosanoid | 1.66E-11 | Increased | 2.397 |
| Core matrisome | Cell movement | 5.4E-09 | Increased | 2.667 |
| | Migration of cells | 1.3E-08 | Increased | 2.257 |

### *APOE* 33 brains: Definite AD vs control

| Enriched pathway | Diseases or Functions annotation | p-value | Predicted Activation State | Activation z-score |
|---|---|---|---|---|
| Matrisome | Migration of cells | 4.5E-29 | Increased | 4.788 |
| | Migration of phagocytes | 1.21E-17 | Increased | 2.629 |
| | Chemotaxis | 5.93E-17 | Increased | 4.423 |
| | Cell movement of myeloid cells | 1.63E-16 | Increased | 3.412 |
| | Cell movement of phagocytes | 2.2E-16 | Increased | 3.585 |
| | Chemoattraction | 2.3E-13 | Increased | 3.068 |
| | Chemotaxis of myeloid cells | 5.43E-12 | Increased | 3.368 |
| | Chemotaxis of phagocytes | 9.48E-12 | Increased | 3.318 |
| | Recruitment of myeloid cells | 2.44E-09 | Increased | 3.082 |
| | Recruitment of phagocytes | 5.98E-09 | Increased | 3.093 |
| | Activation of cells | 2.29E-18 | Increased | 3.882 |
| | Inflammatory response | 1.49E-16 | Increased | 4.27 |
| | Quantity of cells | 9.78E-13 | Increased | 3.676 |
| | Activation of phagocytes | 5.01E-11 | Increased | 3.101 |
| | Activation of myeloid cells | 9.7E-11 | Increased | 3.028 |
| | Stimulation of cells | 1.75E-10 | Increased | 3.876 |
| | Immune response of cells | 3.47E-08 | Increased | 2.574 |
| | Synthesis of eicosanoid | 7.81E-08 | Increased | 2.283 |

### AD brains including *APOE* 44 carriers: Severe vs Mild

| Enriched pathways | Diseases or Functions annotation | p-Value | Predicted Activation State | Activation z-score |
|---|---|---|---|---|
| Matrisome associated | Cell movement of myeloid cells | 1.77E-10 | Increased | 3.152 |
| | Chemotaxis of myeloid cells | 2.50E-08 | Increased | 3.055 |
| | Recruitment of phagocytes | 2.28E-10 | Increased | 2.953 |
| | Cell movement of phagocytes | 1.41E-08 | Increased | 2.918 |
| | Chemotaxis of phagocytes | 3.49E-07 | Increased | 2.73 |
| | Recruitment of myeloid cells | 1.15E-07 | Increased | 2.615 |
| | Activation of cells | 1.66E-13 | Increased | 3.46 |
| | Inflammatory response | 1.38E-10 | Increased | 3.098 |
| | Immune response of cells | 1.56E-08 | Increased | 2.975 |
| | Synthesis of lipid | 4.13E-08 | Increased | 3.285 |
| | Synthesis of fatty acid | 4.10E-06 | Increased | 2.926 |
| | Synthesis of eicosanoid | 1.88E-06 | Increased | 2.759 |
| | Synthesis of prostaglandin | 3.29E-06 | Increased | 2.6 |
| | Synthesis of prostaglandin E2 | 1.07E-05 | Increased | 2.408 |
| | Synthesis of steroid | 5.77E-06 | Increased | 1.547 |
| Matrisome | Cell movement of myeloid cells | 7.5E-12 | Increased | 3.376 |
| | Chemotaxis of myeloid cells | 5.6E-07 | Increased | 3.214 |
| | Cell movement of phagocytes | 3.2E-10 | Increased | 3.172 |
| | Recruitment of phagocytes | 8E-10 | Increased | 2.99 |
| | Chemotaxis of phagocytes | 4.7E-06 | Increased | 2.907 |
| | Recruitment of myeloid cells | 1.9E-07 | Increased | 2.673 |
| | Migration of phagocytes | 3.5E-12 | Increased | 2.315 |
| | Inflammatory response | 3.1E-11 | Increased | 3.562 |
| | Activation of cells | 3E-11 | Increased | 3.557 |
| | Immune response of cells | 6E-08 | Increased | 2.719 |
| | Activation of phagocytes | 2.4E-07 | Increased | 2.36 |
| | Synthesis of fatty acid | 6E-06 | Increased | 3.107 |
| | Synthesis of eicosanoid | 7.6E-07 | Increased | 2.956 |
| | Synthesis of prostaglandin | 4.2E-06 | Increased | 2.277 |
| Core matrisome | Cell movement | 2.9E-05 | Increased | 2.378 |
| | Migration of cells | 0.00015 | Increased | 2.299 |
| | Attachment of cells | 1.6E-08 | Increased | 2.217 |

### Supplemental Data Table 4. Demographics and disease status of the brain bank cohorts.

| **Brain banks** | | | **MSBB** | | **ROSMAP** |
|---|---|---|---|---|---|
| Brain regions | PFC | STG | PHG | IFG | DLPFC |
| Number of *APOE* 44 AD | 9 | 10 | 7 | 8 | 5 |
| Number of *APOE* 33 AD | 46 | 44 | 42 | 45 | 55 |
| Number of *APOE* 33 Control | 15 | 12 | 13 | 11 | 55 |
| Age (years) | 85.2±9.6 | 85.0±9.6 | 85.0±9.7 | 85.3±9.7 | 86.7±4.5 |
| Male (%) | 35.6 | 37.5 | 38.1 | 36.0 | 35.8 |
| Number of CDR = 0 | 35 | 33 | 32 | 27 | 198 |
| Number of CDR = 0.5 | 39 | 33 | 32 | 38 | 167 |
| Number of CDR >= 1 | 187 | 174 | 151 | 157 | 246 |
| Total | 261 | 240 | 215 | 222 | 623 |

Abbreviations: *APOE*, apolipoprotein E; AD, Alzheimer's disease; MSBB, Mount Sinai Brain Bank; ROSMAP, the Religious Orders Study (ROS) and the Memory and Aging Project (MAP); PFC, prefrontal cortex, STG, superior temporal gyrus; PHG, parahippocampal gyrus; IFG, inferior frontal gyrus; DLPFC, dorsolateral prefrontal cortex; CDR, Clinical Dementia Rating.

### Supplemental Data Table 5. Cell type proportion changes by *APOE* genotype or AD phenotype.

| Cell | Region | Comparison | p-value | FDR |
|---|---|---|---|---|
| mic | BA10 | *APOE* 44 AD vs *APOE* 33 AD | 0.272 | 0.612 |
| mic | BA10 | *APOE* 44 AD vs *APOE* 33 Control | 0.223 | 0.582 |
| mic | BA10 | *APOE* 33 AD vs *APOE* 33 Control | 0.586 | 0.823 |
| mic | BA22 | *APOE* 44 AD vs *APOE* 33 AD | 0.049 | 0.330 |
| mic | BA22 | *APOE* 44 AD vs *APOE* 33 Control | 0.018 | 0.221 |
| mic | BA22 | *APOE* 33 AD vs *APOE* 33 Control | 0.306 | 0.612 |
| mic | BA36 | *APOE* 44 AD vs *APOE* 33 AD | 0.604 | 0.823 |
| mic | BA36 | *APOE* 44 AD vs *APOE* 33 Control | 0.033 | 0.288 |
| mic | BA36 | *APOE* 33 AD vs *APOE* 33 Control | 0.013 | 0.199 |
| mic | BA44 | *APOE* 44 AD vs *APOE* 33 AD | 0.198 | 0.566 |
| mic | BA44 | *APOE* 44 AD vs *APOE* 33 Control | 0.116 | 0.439 |
| mic | BA44 | *APOE* 33 AD vs *APOE* 33 Control | 0.393 | 0.674 |

| Cell | Region | Comparison | p-value | FDR |
|---|---|---|---|---|
| neu | BA10 | *APOE* 44 AD vs *APOE* 33 AD | 0.944 | 0.973 |
| neu | BA10 | *APOE* 44 AD vs *APOE* 33 Control | 0.265 | 0.612 |
| neu | BA10 | *APOE* 33 AD vs *APOE* 33 Control | 0.219 | 0.582 |
| neu | BA22 | *APOE* 44 AD vs *APOE* 33 AD | 0.814 | 0.921 |
| neu | BA22 | *APOE* 44 AD vs *APOE* 33 Control | 0.160 | 0.483 |
| neu | BA22 | *APOE* 33 AD vs *APOE* 33 Control | 0.041* | 0.311 |
| neu | BA36 | *APOE* 44 AD vs *APOE* 33 AD | 0.441 | 0.679 |
| neu | BA36 | *APOE* 44 AD vs *APOE* 33 Control | 0.111 | 0.439 |
| neu | BA36 | *APOE* 33 AD vs *APOE* 33 Control | 0.006* | 0.148 |
| neu | BA44 | *APOE* 4 AD vs *APOE* 33 AD | 0.441 | 0.679 |
| neu | BA44 | *APOE* 44 AD vs *APOE* 33 Control | 0.957 | 0.973 |
| neu | BA44 | *APOE* 33 AD vs *APOE* 33 Control | 0.644 | 0.829 |

| Cell | Region | Comparison | p-value | FDR |
|---|---|---|---|---|
| ast | BA10 | *APOE* 44 AD vs *APOE* 33 AD | 0.694 | 0.852 |
| ast | BA10 | *APOE* 44 AD vs *APOE* 33 Control | 0.288 | 0.612 |
| ast | BA10 | *APOE* 33 AD vs *APOE* 33 Control | 0.422 | 0.679 |
| ast | BA22 | *APOE* 44 AD vs *APOE* 33 AD | 0.088 | 0.438 |
| ast | BA22 | APOE44 AD vs *APOE* 33 Control | 0.466 | 0.699 |
| ast | BA22 | *APOE* 33 AD vs APOE33 Control | 0.696 | 0.852 |
| ast | BA36 | *APOE* 44 AD vs *APOE* 33 AD | 0.026 | 0.260 |
| ast | BA36 | *APOE* 44 AD vs *APOE* 33 Control | 0.347 | 0.632 |
| ast | BA36 | *APOE* 33 AD vs *APOE* 33 Control | 0.093 | 0.438 |
| ast | BA44 | *APOE* 44 AD vs *APOE* 33 AD | 0.121 | 0.439 |
| ast | BA44 | *APOE* 44 AD vs *APOE* 33 Control | 0.161 | 0.483 |
| ast | BA44 | *APOE* 33 AD vs *APOE* 33 Control | 0.417 | 0.679 |

| Cell | Region | Comparison | p-value | FDR |
|---|---|---|---|---|
| end | BA10 | *APOE* 44 AD vs *APOE* 33 AD | 0.979 | 0.979 |
| end | BA10 | *APOE* 44 AD vs *APOE* 33 Control | 0.378 | 0.667 |
| end | BA10 | *APOE* 33 AD vs *APOE* 33 Control | 0.302 | 0.612 |
| end | BA22 | *APOE* 44 AD vs *APOE* 33 AD | 0.900 | 0.948 |
| end | BA22 | *APOE* 44 AD vs *APOE* 33 Control | 0.007 | 0.148 |
| end | BA22 | *APOE* 33 AD vs *APOE* 33 Control | 0.003 | 0.148 |
| end | BA36 | *APOE* 44 AD vs *APOE* 33 AD | 0.732 | 0.861 |
| end | BA36 | *APOE* 44 AD vs *APOE* 33 Control | 0.347 | 0.632 |
| end | BA36 | *APOE* 33 AD vs *APOE* 33 Control | 0.278 | 0.612 |
| end | BA44 | *APOE* 44 AD vs *APOE* 33 AD | 0.145 | 0.483 |
| end | BA44 | *APOE* 44 AD vs *APOE* 33 Control | 0.124 | 0.439 |
| end | BA44 | *APOE* 33 AD vs *APOE* 33 Control | 0.649 | 0.829 |

| Cell | Region | Comparison | p-value | FDR |
|---|---|---|---|---|
| oli | BA10 | *APOE* 44 AD vs *APOE* 33 AD | 0.852 | 0.929 |
| oli | BA10 | *APOE* 44 AD vs *APOE* 33 Control | 0.532 | 0.779 |
| oli | BA10 | *APOE* 33 AD vs *APOE* 33 Control | 0.618 | 0.825 |
| oli | BA22 | *APOE* 44 AD vs *APOE* 33 AD | 0.339 | 0.632 |
| oli | BA22 | *APOE* 44 AD vs *APOE* 33 Control | 0.598 | 0.823 |
| oli | BA22 | *APOE* 33 AD vs *APOE* 33 Control | 0.233 | 0.582 |
| oli | BA36 | *APOE* 44 AD vs *APOE* 33 AD | 0.094 | 0.438 |
| oli | BA36 | *APOE* 44 AD vs *APOE* 33 Control | 0.069 | 0.415 |
| oli | BA36 | *APOE* 33 AD vs *APOE* 33 Control | 0.720 | 0.861 |
| oli | BA44 | *APOE* 44 AD vs *APOE* 33 AD | 0.882 | 0.945 |
| oli | BA44 | *APOE* 44 AD vs *APOE* 33 Control | 0.847 | 0.929 |
| oli | BA44 | *APOE* 33 AD vs *APOE* 33 Control | 0.749 | 0.865 |

mic, microglia; neu, neurons; ast, astrocytes; end, endothelial cells; oli, oligodendrocytes

### Supplemental Data Table 6. Genetic identity-by-descent estimation of genome-edited isogenic *APOE* hiPSC lines to original source fibroblasts

| Fibroblast/Individual ID | *APOE* genotype | Isogenic *APOE* hiPSC line ID | PI HAT |
|---|---|---|---|
| **8** | 33 | TCW1E33-1C6 | 0.9999 |
| | 33 | TCW1E33-1C8 | 0.9999 |
| | 33 | TCW1E33-1F1 | 1 |
| | 44 | TCW1E44-2C2 | 1 |
| | 44 | TCW1E44-2E3 | 1 |
| | 44 | TCW1E44-1D1 | 0.9999 |
| **10** | 33 | TCW2E33-2E3 | 1 |
| | 33 | TCW2E33-3A5 | 0.9999 |
| | 33 | TCW2E33-3D11 | 0.9999 |
| | 44 | TCW2E44-4B1 | 1 |
| | 44 | TCW2E44-4B4 | 1 |
| | 44 | TCW2E44-4B12 | 1 |

**Supplemental Data Figure 1. Raw blots used in main figures.**

**Fig. 5f**

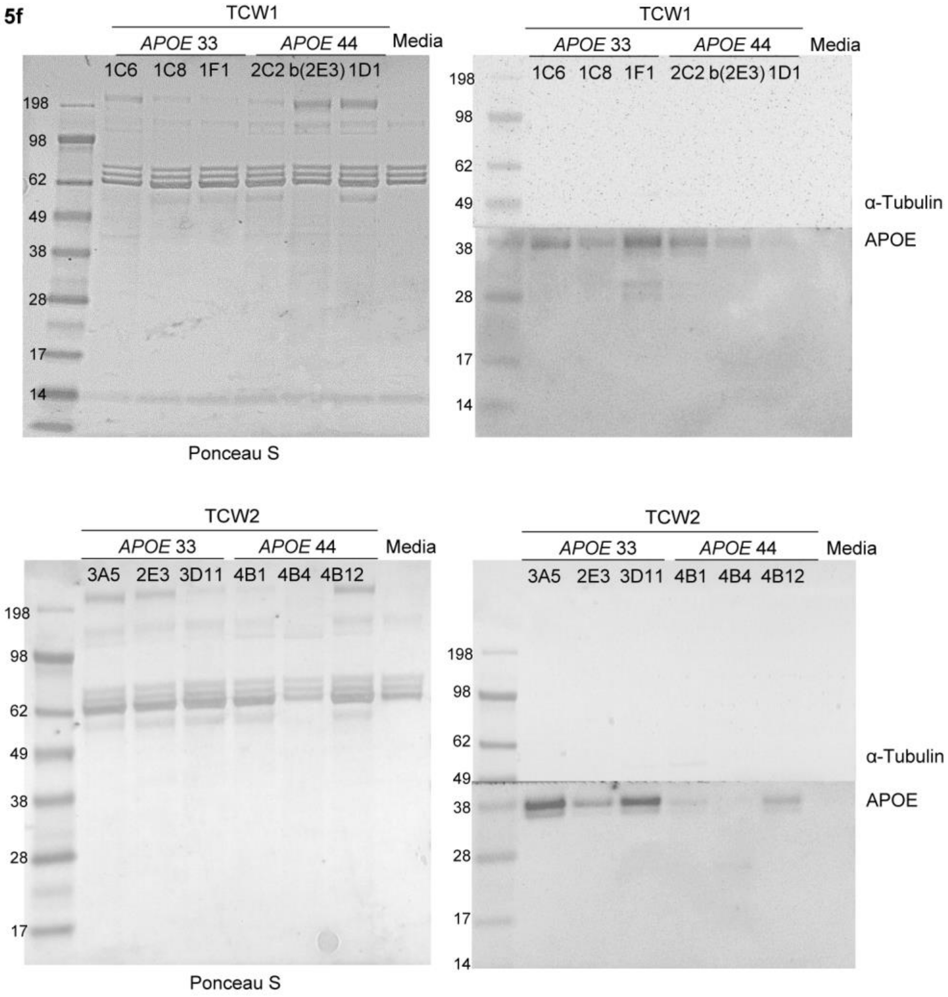
